# The ‘squalene route’ to carotenoid biosynthesis is widespread in *Bacteria*

**DOI:** 10.1101/2021.12.22.473825

**Authors:** Carlos Santana-Molina, Valentina Henriques, Damaso Hornero-Méndez, Damien P. Devos, Elena Rivas-Marin

**Author notes:** Addresses for correspondence to CSM, DPD and ERM.

## Abstract

Squalene is mostly associated with the biosynthesis of polycyclic triterpenes. Although there have been suggestions that squalene could be involved in the biosynthesis of carotenoids, functionally and evolutionarily related to polycyclic triterpenes, evidence of this ‘squalene route’ in nature was lacking. We demonstrate that planctomycetes synthesize C30 carotenoids via squalene and that this ‘squalene route’ is widely distributed in *Bacteria*. We also investigated the functional roles of hopanoids and carotenoids in *Planctomycetes* and show that their protective functions under stress conditions are complementary. Our evolutionary analyses suggest that the C30 carotenoid biosynthetic pathway is the most ancestral, with a potential origin in *Firmicutes* or *Planctomycetes*. In addition, we propose an evolutionary scenario to explain the diversification of the different carotenoid and squalene pathways. Together, these results improve the evolutionary contextualization of these molecules. Likewise, the widespread occurrence of the squalene route in bacteria increases the functional repertoire of squalene.

## Introduction

Carotenoids are isoprenoid lipids found in all photosynthetic and some non-photosynthetic organisms. The most abundant carotenoids are produced by photosynthetic organisms and have C40 backbones, although some chemoorganotrophic bacteria are capable of producing *de novo* C30, C45 or C50 carotenoids (1, 2). Carotenoid biosynthesis is evolutionarily related to the biosynthesis of polycyclic triterpenes, such as hopanoids and sterols, as the enzymes involved are homologues (3, 4). Squalene (C_30_H_50_) is the precursor of polycyclic triterpenes, which can be synthesized via two routes: the HpnCDE pathway, found mostly in bacteria; and squalene synthase (Sqs), a single enzyme found in the three domains of life. HpnC, HpnD and Sqs belong to the trans-isoprenyl diphosphate synthases head-to-head (Trans IPPS HH) family, to which the enzymes initiating C30 and C40 carotenoid biosynthesis also belong (Fig. S1). These Trans IPPS HHs generate the initial backbones of polycyclic triterpenes or carotenoids, which then become the substrate for specific amino oxidases (also known as phytoene desaturases): CrtN/P and HpnE act on C30 backbones, and CrtI/D or CrtP-Q/CrtH act on C40 backbones (Fig. S1).

The production of C30 carotenoids is widespread in *Bacteria*, but how the precursor is synthesized is unclear in most cases; thus far, bacterial C30 carotenoid production has been characterized only via CrtM, which is specific to *Firmicutes*. Some bacteria, such as planctomycetes, have C30-specific amino oxidases and subsequent carotenoid-modifying enzymes, but usually no Trans IPPS HHs associated with carotenoid precursor synthesis. Instead, planctomycetal genomes encode the HpnCDE enzymes responsible for squalene production, suggesting that squalene could be the substrate for these C30-specific amino oxidases (4). This possibility is supported by three observations. First, squalene can act as the substrate for C30- but not C40- specific amino oxidases, however, this ‘squalene route’ has only been artificially demonstrated so far (5). Second, a sterol synthesis-deficient mutant of the planctomycete *Gemmata obscuriglobus* that accumulates squalene shows a brighter red pigmentation (6). And third, interrupting the *hpnE* genes in the planctomycete *Planctopirus limnophila* and in the alphaproteobacterium *Methylobacterium extorquens* results in non-pigmented colonies due to the lack of carotenoid production (4, 7).

Carotenoids are also expected to be functionally related to polycyclic triterpenes. Carotenoids have important biological properties, including photoprotection and modification of the fluidity, permeability and stability of the cellular membranes (8–10). However, their functional similarities to and differences from polycyclic triterpenes are unclear.

Here, we investigate the synthesis of carotenoids in the planctomycete *P. limnophila*. We demonstrate the existence of the squalene route to carotenoid biosynthesis in nature and further report its widespread occurrence in *Bacteria*. We also investigate the potential roles of hopanoids and carotenoids in *Planctomycetes*, showing for the first time that these molecules have complementarity protective function in these bacteria. These novel insights, together with our evolutionary analyses, contextualize the ancestral diversification of terpenoid metabolism comprising polycyclic triterpenes and carotenoids. This report of the widespread occurrence of the squalene route in *Bacteria* decouples squalene from the biosynthesis of polycyclic triterpenes, to which it has traditionally been associated, and increases its functional repertoire.

## Results

### Planctomycetes produce C30 carotenoids using squalene as a precursor

To decipher carotenoid production in planctomycetes, we performed random mutagenesis by Tn5 transposition to select *P. limnophila* colonies with altered pigmentation. The transposition events in the selected clones mapped to the genes *hpnD, hpnE, crtN, crtP, aldH, crtQ* and *crtO*, which comprised all genes previously identified computationally with the exception of *hpnC* (4). We deleted *hpnC* by directed mutagenesis. The *hpnC, hpnD, hpnE* and *crtN* mutants, resulted in colorless colonies, while the other mutants showed colonies with altered pigmented, with colors ranging from light yellow to bright red (Fig. 1a).

**Figure 1:**
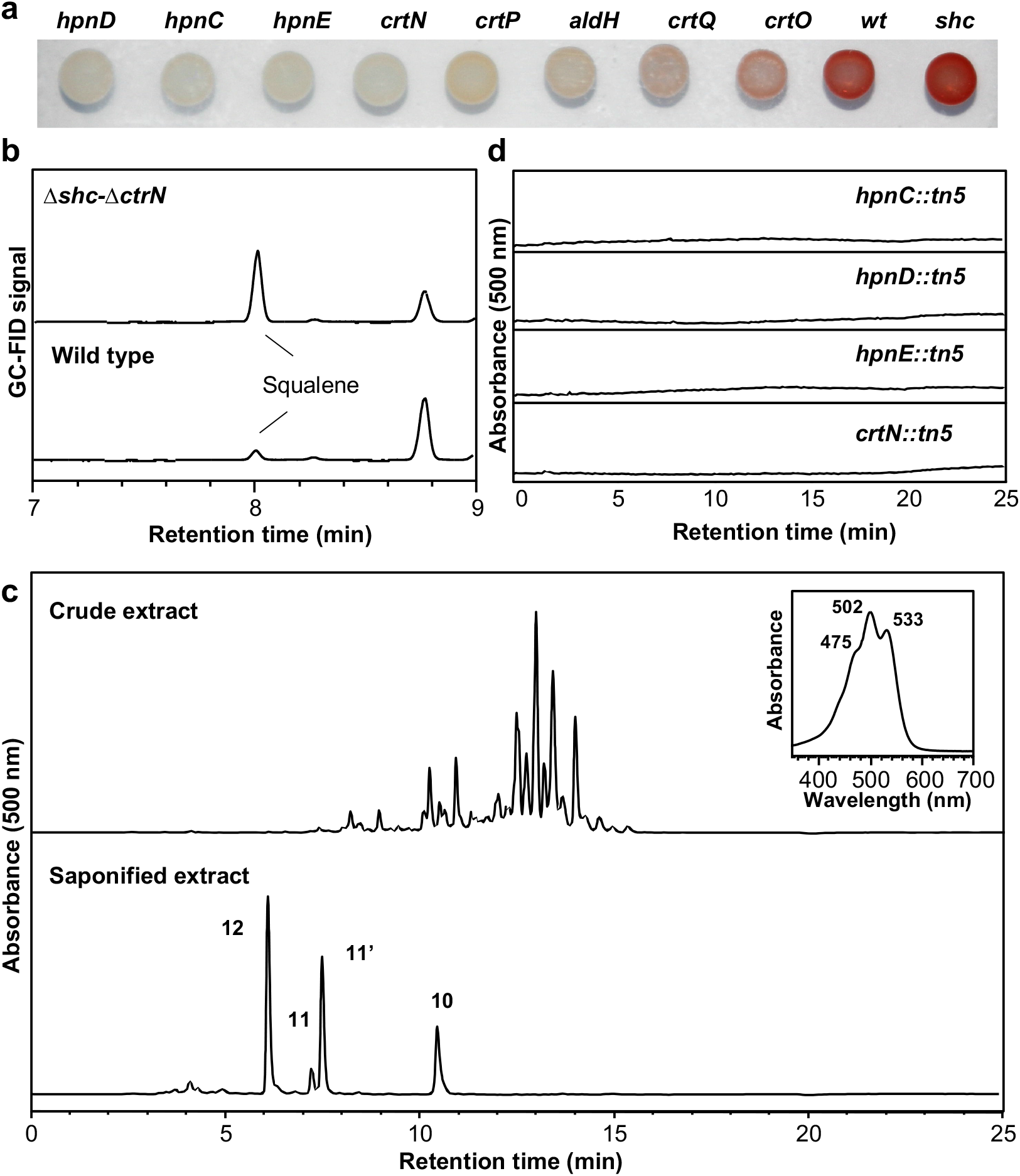
*P. limnophila* carotenoids analysis. a) Culture drops of wild type and various mutants lacking the indicated genes, b) GC-FID analysis of squalene in wild type and Δ*crtN*-Δ*shc* mutant strains, c) HPLC separation of carotenoid pigments present in crude and saponified (alkaline hydrolysis) extracts obtained from wild type strain and total UV-visible spectrum for the crude extract. Peak numbers are in accordance with the chromatograms in Figure 2 and the pathway scheme (Fig. 3): 4,4’-diapolycopenoic acid (10); 4,4’-diapolycopen-4’-al-4-oic acid (11 & 11’); 4,4’-diapolycopen-4,4’-dioic acid (12). d) HPLC chromatograms corresponding to the carotenoid analysis for transposon-inserted mutants (*hpnC, hpnD, hpnE* and *crtN*). HPLC detection wavelength at 500 nm.

We also constructed a hopanoid-deficient mutant by deleting the *shc* (squalene hopene cyclase) gene. The resulting colonies grew under standard conditions, thus discounting the essentiality of hopanoids in *P. limnophila*, in contrast to the essentiality of sterols in its close relative, *G. obscuriglobus* (6). The mutant colonies displayed an intense red color (Fig. 1a), indicating that the accumulated squalene (or its precursors) is re-directed towards carotenoid production, as observed in *M. extorquens* (7, 11). Indeed, we confirmed squalene production via HpnCDE using the Δ*crtN*-Δ*shc* strain, since squalene is the precursor of hopanoids. The production and accumulation of squalene in this mutant was confirmed by gas chromatography with flame-ionization detection (GC-FID) (Fig. 1b). Together with the lack of color in the *hpnCDE* mutants (Fig. 1a), these results suggest a role for squalene as an intermediate in carotenoid biosynthesis.

To characterize the carotenoid pigments produced by wild type *P. limnophila* and the selected mutants, the corresponding cell extracts were analyzed by high-performance liquid chromatography (HPLC), and UV-visible spectra were obtained for each peak. Wild type *P. limnophila* showed a complex HPLC profile with many peaks having UV-visible spectra in agreement with chromophore structures that have 11 to 13 conjugated double bonds (Fig. 1c). The fact that most peaks presented almost identical UV-visible spectra but different chromatographic mobility suggested the possibility that these peaks corresponded to an array of different esterified forms. This was analyzed upon alkaline hydrolysis of wild type extract, which produced a simpler chromatogram (Fig. 1c), confirming the esterified nature of the pigments. Moreover, acidification of the hydrolyzed pigments solution was necessary to transfer them to organic solvent (diethyl ether), suggesting that the major pigments in the wild type were acidic carotenoids in which the native form was glycosyl esters, in agreement with previous studies (12). In fact, the spectroscopic (UV-visible and MS) and chromatographic properties of the major peaks in the saponified extract from wild type were in agreement with that of carotenoid acids derived from 4,4’-diapolycopene, a C30 carotenoid observed in other bacterial species such as *Methylobacterium rhodium* (formerly *Pseudomonas rhodos*) (13, 14), *Rubritella squalenifaciens* (15, 16), *Planococcus maritimus* (17) and *Bacillus firmus* (18). In addition, we did not find C40 carotenoids – such as lycopene, neurosporene and β-carotene – in wild type or mutants. Taking these data together, we thus tentatively identified *P. limnophila* pigments as C30 carotenoids, the most predominant ones were carotenoid acids derived from 4,4’-diapolycopene (Fig. 1c). We note that carotenoids have been detected in the planctomycetes *Rhodopirellula rubra* LF2^T^ and *Rubinisphaera brasiliensis* Gr7 (19). However, the nature and biosynthetic pathways of these carotenoids are still unclear.

To explore the action of the detected enzymes, pigment extracts from different mutants were analyzed by HPLC-DAD (Fig. 2 and Fig. S2). Additionally, the mass spectra for the most predominant compounds were obtained by HPLC-MS (APCI) (Fig. S3). The *hpnC, hpnD* and *hpnE* transposon mutants lacked carotenoids (Fig. 1d). Similarly, carotenoids were absent in the *crtN* mutant (Fig. 1b). The *crtP::tn5* mutant showed light orange pigmentation and accumulated 4,4’-diapolycopene (1) as the most predominant pigment (Fig. 2). The UV-visible of this compound (Fig. S2) confirmed the presence of eleven conjugated double bonds in its structure, and the mass spectrum (Fig. S3) showed a prominent ion corresponding to the protonated molecule [M+H]^+^ at 401.31 which is consistent with the formula C_30_H_40_ (Mw=400.31) of 4,4’-diapolycopene. All the pathway precursors, namely 4,4’-diaponeurosporene (2), 4,4’-diapo-ζ-carotene (3) and 4,4’-diapophytofluene (4), leading to the formation of 4,4’-diapolycopene by the action of CrtN (4,4’-diapophytoene desaturase) were also detected. The UV-visible spectra for these compounds (Fig. S2) agreed with the extension of the conjugated double bond system of the proposed structures (Fig. 3). The *aldH::tn5* mutant (interrupted in the gene coding for a carotenoid aldehyde dehydrogenase) showed red pigmentation and presented the expected carotenoids containing aldehyde end-groups such as 4,4’-diapolycopen-4-al (5) and 4,4’-diapolycopene-dial (7). Both, UV-visible and MS spectra (Fig. S2 and S3) were consistent with the proposed C30 structures. Interestingly, this mutant also accumulated other pigments that we provisionally identified, based on their UV-visible spectra and chromatographic properties, as hydroxy derivatives of 4,4’-diapolycopene: 4,4’-diapolycopene-4-ol (6) and 4,4’-diapolycopene-4,4’-diol (9). These two pigments presented the same UV-visible spectrum as that of 4,4’-diapolycopene (Fig. S2) but had higher polarity; hydroxylation is the only possible structural modification that would increase the polarity of the derivatives without modifying the chromophore properties. We also tentatively identified a carotenoid containing one aldehyde group and one hydroxy group: 4,4’-diapolycopene-4-ol-4’-al (8). The occurrence of hydroxylated diapolycopene derivatives raises the question of whether the introduction of hydroxy group is due to an additional hydroxylase activity of CrtP, to the action of an unknown hydroxylase enzyme, or even to a spontaneous step(20). The *crtQ::tn5* mutant accumulated carotenoic acids derived from the action of the carotenoid aldehyde dehydrogenase (*aldH*), namely 4,4’-diapolycopenoic acid (10), 4,4’-diapolycopen-4’-al-4-oic acid (11 & 11’) and 4,4’-diapolycopen-4,4’-dioic acid (12). As shown before, these were also the major carotenoids found in the wild type extract. The structure for these C30 compounds were in accordance with their UV-visible and MS spectra (S 2 and S3). It is interesting to note that the carboxylic acid moiety can be formed not only by oxidation of an intermediary aldehyde (via AldH) but also by a putative reaction introducing a keto group into a carbon with a preexisting hydroxy group. In this way, both aldehyde and hydroxy 4,4’-diapolycopene derivatives would be transformed into the same carotenoic acid compounds. The *crtO::tn5* mutant contained pigments (peaks 13, 14, 15 and 16) with UV-visible spectra similar to those of 4,4’-diapolycopenoic acids but with increased polarities, which were consistent with the acylation with sugar moieties to produce carotenoic acid glycosyl esters. The additional acylation of the sugar derivatives with different fatty acids by the action of CrtO (acyl transferase) produced the complex pigment profile observed in the wild type strain (Fig. 1d). Additional experimental work is needed to identify both the sugar moieties and the fatty acids involved in the formation of the carotenoid acid glycosyl derivatives responsible for the native carotenoid profile and color of *P. limnophila*.

**Figure 2:**
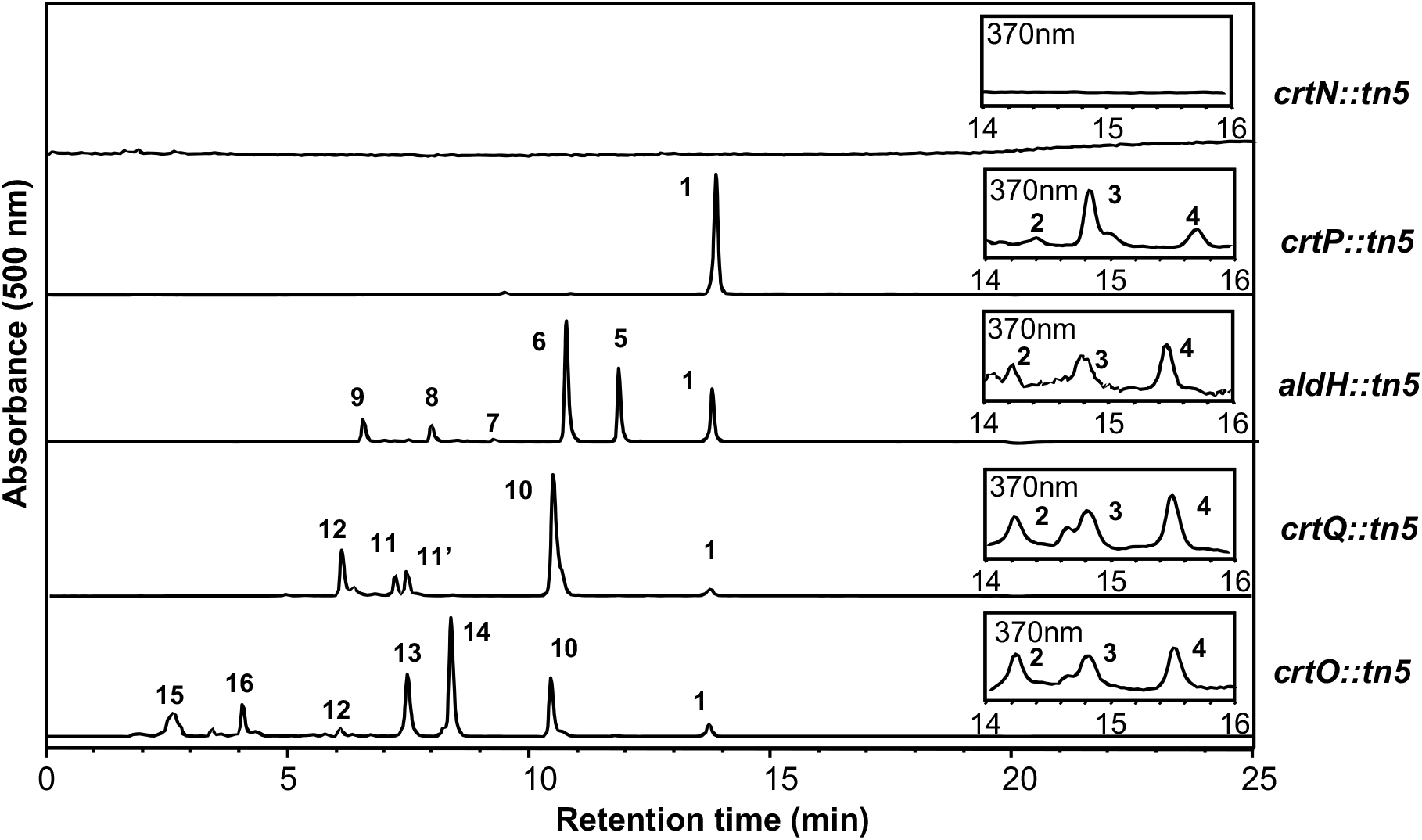
HPLC chromatograms corresponding to the analyses of *P. limnophila* carotenoid mutants. Detection wavelengths at 370 and 500 nm. Peaks: 4,4’-diapolycopene (1); 4,4’-diaponeurosporene (2); 4,4’-diapo-ζ-carotene (3); 4,4’-diapophytofluene (4); 4,4’-diapolycopen-4-al (5); 4,4’-diapolycopene-4-ol (6); 4,4’-diapolycopene dial (7); 4,4’-diapolycopene-4-ol-4’-al (8); 4,4’-diapolycopene-4,4’-diol (9); 4,4’-diapolycopenoic acid (10); 4,4’-diapolycopen-4’-al-4-oic acid (11 & 11’); 4,4’-diapolycopen-4,4’-dioic acid (12); glycosyl esters of 4,4’-diapolycopenoic acid (13 & 14); and glycosyl esters of 4,4’-diapolycopen-4,4’-dioic acid (15 & 16).

**Figure 3:**
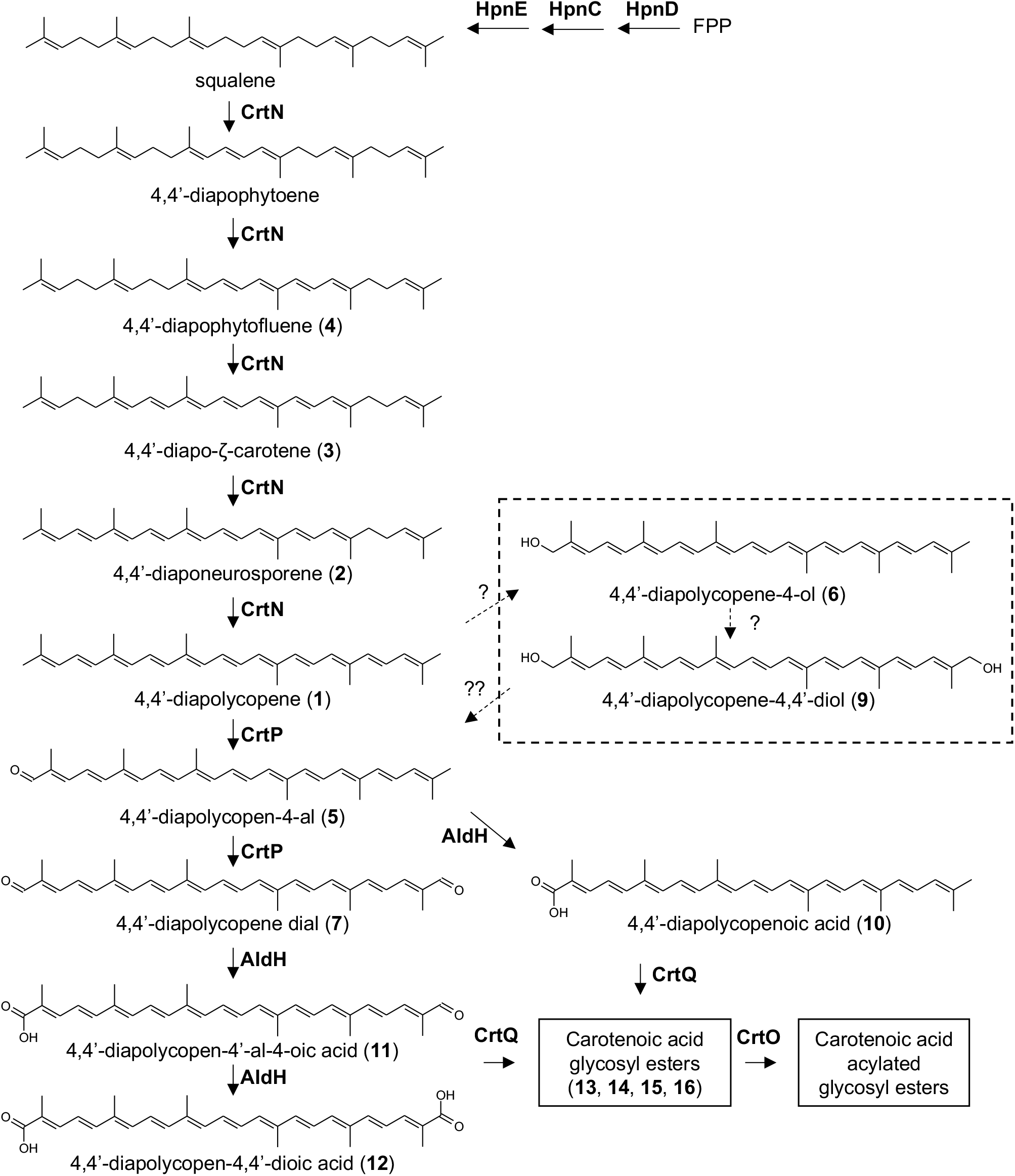
Proposed pathway for carotenoid biosynthesis in *P. limnophila*. Compound numbers are in accordance with peak numbers in Figure 2.

### Heterologous reconstruction in *Escherichia coli*

To corroborate the synthesis of carotenoids from squalene in *P. limnophila*, we assembled a heterologous expression system in *E. coli* BL21, a strain that lacks both carotenoids and hopanoids. *E. coli* was simultaneously transformed with three plasmids (Table S3). The first plasmid ensured isopentenyl diphosphate (IPP) production, to enhance the biosynthesis of isoprenoid derivatives. The second plasmid carried the squalene synthesis genes (*hpnCDE* or *sqs*). Different versions of the third plasmid contained the downstream carotenoid modification genes in an additive fashion (*crtN, crtP, aldH, crtO* and *crtQ*). In addition to these strains, alternative versions of plasmid 2 were also constructed and used as controls. One version contained only the *hpnC* and *hpnD* genes. Colonies containing this plasmid were colorless, confirming that carotenoids were produced via squalene and not via intermediates of the HpnCDE pathway (Fig. S4). Another version contained the cyanobacterial *sqs* gene for squalene production. An *E. coli* strain containing this plasmid yielded the same carotenoids as the strain with the plasmid bearing the *hpnCDE* genes, although with different proportions, which confirmed that both pathways are equivalent for carotenoid synthesis (Fig. S4). HPLC analysis verified that *E. coli* expressing the whole pathway produced a mixture of carotenoids, including 4,4’-diapolycopene (the most abundant), 4,4’-diapolycopen-4-al, 4,4’-diapolycopen-4’-al-4-oic acid, 4,4’-diapolycopen-4,4’-dioic acid and 4,4’-diapolycopenoic acid. The glycosyl- and ester-modified carotenoids did not appear in the *E. coli* extracts, unlike in the *P. limnophila* extracts, although similar precursors were observed (Fig. 2 and Fig. S4).

Compiling the results from genetic, bioinformatic and carotenoid analyses, we propose a tentative pathway for the synthesis of C30 carotenoids in *P. limnophila* via the squalene route (Fig. 3).

### Functional characterization of C30 carotenoids and hopanoids in *P. limnophila*

To study the role of each of the triterpenoids, we constructed deletion mutants of some of the genes previously inserted by random mutagenesis. We selected the following *P. limnophila* strains to study the functional role of each terpenoid in isolation: *P. limnophila* Δ*shc* and Δ*crtN*, which do not produce hopanoids or carotenoids, respectively; Δ*crtN*-Δ*shc*, which produces squalene but not hopanoids or carotenoids; and Δ*hpnD*, which is unable to produce any of these molecules. We observed no statistically significant difference in growth rate between the mutants and the wild type, even when they were grown at different temperatures (Fig. 4a; Fig. S5a). However, growth of the Δ*hpnD* and the Δ*crtN*-Δ*shc* mutant strains was slower when the strain was grown on solid plates with 1.5 % agar the absence of stress (Fig. S5b). In the case of the Δ*crtN*-Δ*shc* mutant the wild type growth was recovered when grown at 1 % agar (Fig. S5b). To establish the physiological roles of each molecule, we analyzed the mutants under different stress conditions, including desiccation, osmotic stress and oxidative stress (Fig. 4b-c). We found that *P. limnophila* carotenoids or hopanoids are not linked to any specific protection against a particular stress. However, we observed a general cumulative effect: all triterpenoids are associated with protection against the tested stresses in an incremental fashion. Squalene is slightly protective, and carotenoids and hopanoids have additive protective functions. However, in the case of carotenoids and hopanoids, we cannot rule out the possibility that the elimination of one molecule by interrupting one pathway was compensated by the overproduction of the other molecule. This possibility is supported by the brighter red pigmentation observed in the *G. obscuriglobus* and *P. limnophila* mutant strains lacking sterol and hopanoid, respectively.

**Figure 4:**
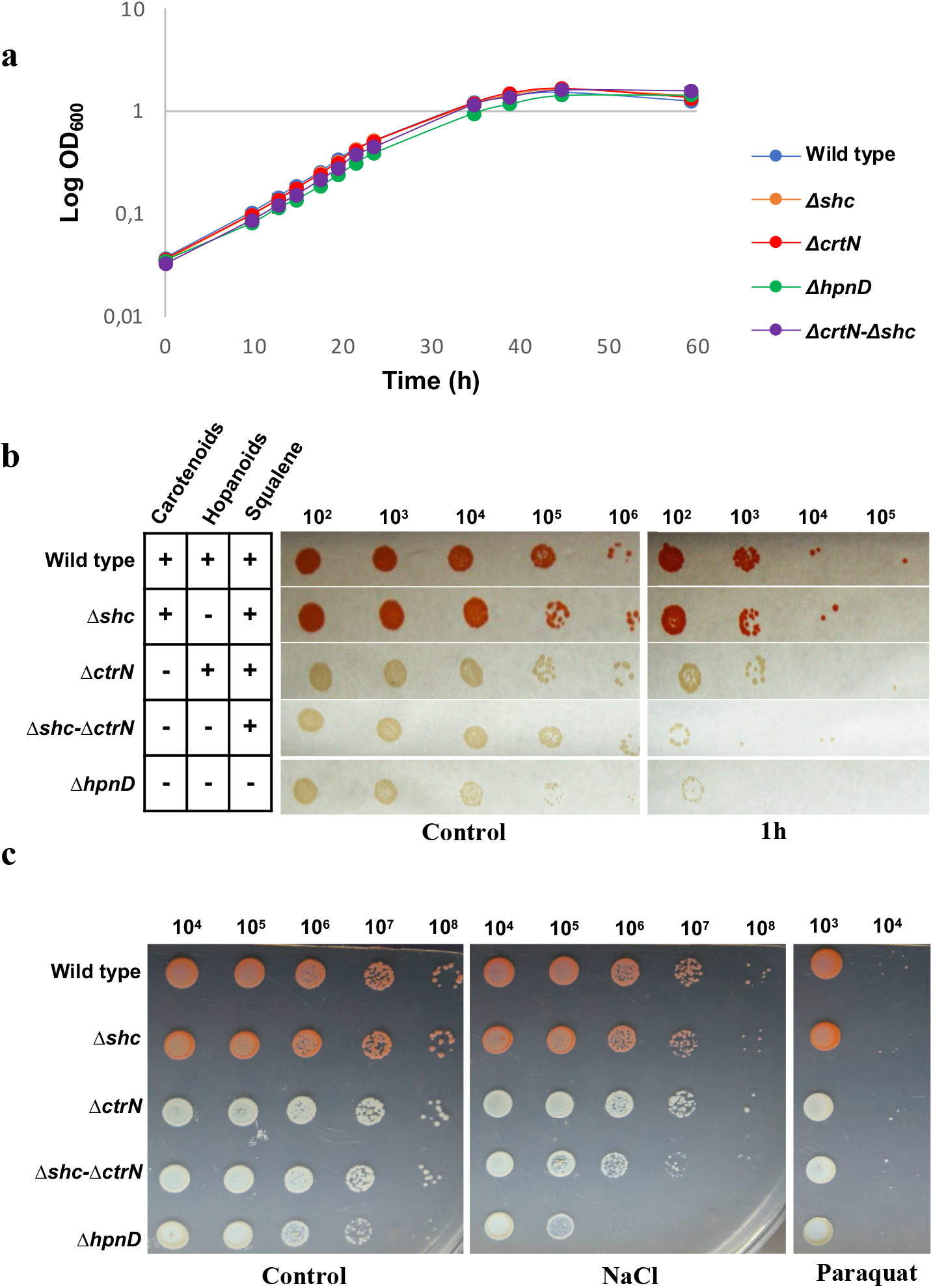
Phenotypic assays of selected *P. limnophila* mutants. a) growth curves, b) growth under desiccation stress (1h), and c) growth in plates (1 % agar) under osmotic (15 mM NaCl) and oxidative stress (2 μM paraquat).

Finally, the different mutant strains, together with the wild type, were exposed to several freeze/thaw cycles, as carotenoids have been shown to be protective against ice-induced membrane defects (10). The viability of neither the mutants nor the wild type was affected, even after three cycles of freeze/thaw (data not shown).

### Evolution of the C30 carotenoid pathway

We next reconstructed the evolutionary history of the main enzymes that define the carotenoid biosynthetic pathways, the amino oxidases. The inferred phylogenies of CrtN and CrtP provided congruent topologies, which suggests that these enzymes have a common evolutionary history (Fig. 5). This phylogenetic congruency is in agreement with the fact that these genes form an operon together with other carotenoid-related genes, such as *aldH, crtQ* or *crtO*, in most genomes (Fig. S6). CrtN is more widespread than CrtP, which could be related to the fact that it is the first amino oxidase enzyme in the pathway, responsible for joining the two C15 subunits into the C30 backbone. The CrtN/P phylogeny shows a taxonomically mixed topology characterized by *Firmicutes-Bacilli* (and *Clostridia*, which contain only CrtN) branching paraphyletically and basally in the respective subfamilies. Embedded in this group are other bacterial orders such as *Verrucomicrobiales*, *Acidobacteriales* and *Methylococcales*, among others. Another main basal branch in the CrtN/P phylogeny is composed of the *Planctomycetes* phylum (classes *Planctomycetia* and *Phycisphaerae*), which contains other taxonomic groups embedded paraphyletically. These groups encompass bacterial orders with few representatives, such as *Rhizobiales, Acetobacteriales, Rhodobacterales*, *Acidobacteriales* and the euryarchaeal class *Candidatus* Poseidoniia, among others, which contain only CrtN (including MGII archaea from *Candidatus* Thermoplasmatota). This taxonomic and phylogenetic distribution of the C30-specific amino oxidases suggests that there have been multiple lateral gene transfer events between prokaryotes, mainly from *Firmicutes-Bacilli* and *Planctomycetes*.

**Figure 5:**
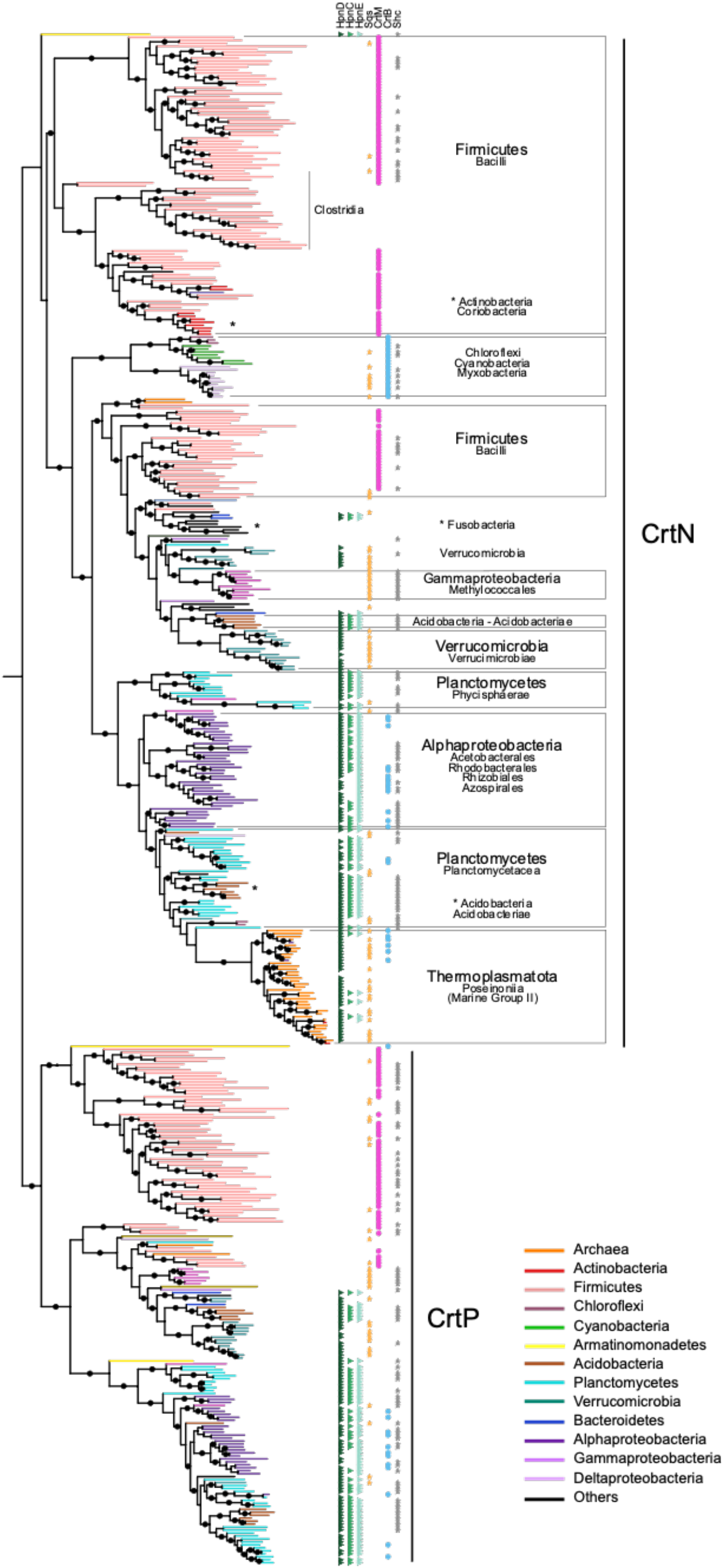
Phylogeny of C30-specific amino oxidases CrtN and CrtP and co-occurrence with genes of interest. The dataset was reduced by phyla by progressive redundancy. Branches are colored according to the prokaryotic phyla. Black circles indicate bootstraps >90 %. Phyla and classes are indicated. The phylogenetic profile includes the presence of HpnCDE (green triangles), Sqs (orange star), CrtM (pink circle), CrtB (blue circle) and Shc (gray star). More information for the taxonomic profile in shown in Data S1.

We next analyzed the potential source of the precursor of these carotenoid amino oxidases in the different organisms. We found that the biosynthesis of 4,4’-diapophytoene by CrtM is restricted to *Firmicutes*, while the squalene pathway (via HpnCDE or Sqs) is found in other organisms bearing the CrtN/P proteins (Fig. 5). This result shows that the C30-specific amino oxidases are usually associated with the presence of 4,4’-diapophytoene (in *Firmicutes*) or squalene biosynthesis, in agreement with our demonstration of C30 carotenoid production from squalene. This is further supported by the presence of *hpnD* in the genomic context of *crtN* of the archaeal class *Ca*. Poseidoniia or of *hpnCDE* in some alphaproteobacterial orders (Fig. S6; Data S2). Thus, our results show that the squalene route to carotenoid production is widespread in *Bacteria* and that the C30 carotenoid pathway has been transferred multiple times. In addition, the biosynthesis of the precursor (4,4’-diapophytoene or squalene) has shifted between different groups.

### Origins of carotenoid pathways

The taxonomic distribution of the genes involved in C30 carotenoid synthesis is more limited than that of squalene or hopanoids (4). This limited distribution narrows the possible taxonomic origin of C30 carotenoid pathways. There are two main pieces of evidence that suggest *Firmicutes* could be at the origin of C30 carotenoids. One is that *Firmicutes* forms the most basal clades in the phylogeny of CrtN and CrtP (Fig. 5), which in this case could be sign of ancestrality or of more intense divergence. The second piece of evidence is that Firmicutes are the only bacteria bearing CrtM, which suggests that the biosynthesis of C30 carotenoids via 4,4’-diapophytoene originated in these organisms. Another interesting fact is the exclusive presence of CrtN in *Clostridia*, which represents an ancient anaerobic class of *Firmicutes* and thus, perhaps, an early origin of the CrtN subfamily. By contrast, the biosynthesis of C30 carotenoids via squalene shows an ancestral evolution in *Planctomycetes* and represents an alternative to the origin of C30 carotenoids that is independent of CrtM (4,4’-diapophytoene synthase).

The carotenoid amino oxidases usually work in pairs, and their evolution provides a broad view of how the different carotenoid pathways evolved. HpnE and CrtN/P act on C30 backbones, while CrtI/D and CrtP-Q/H act in the two main pathways for C40 carotenoid biosynthesis (Fig. S1). One of these pathways is present exclusively in aerobic *Cyanobacteria* and green sulfur anaerobic bacteria (*Chlorobi*), via CrtP-Q and CrtH (Fig. S7). CrtH is phylogenetically distant from the other families, while CrtP-Q from *Cyanobacteria* is closely related to HpnE. Thus, this C40 CrtP-Q/H pathway most likely originated in *Cyanobacteria* or *Chlorobi* with an origin related to HpnE and independent from the other C40 pathway. The other C40 pathway, via CrtI/D, most likely had an ancestral evolution in *Proteobacteria, Bacteroidetes, Actinobacteria* and *Deinococcus*, among others, suggesting lateral gene transfer events between the ancestor of these phyla (Fig. S7). While CrtD is more conserved, CrtI has a limited distribution that is restricted mainly to *Proteobacteria* and *Actinobacteria*. However, *Bacteroidetes*, for example, has a different carotenoid amino oxidase for C40 carotenoid biosynthesis, CrtDb, instead of the ‘classical’ CrtI. This CrtDb has been transferred by lateral gene transfer between the ancestor of *Bacteroidetes*, *Actinobacteria*, *Thermoplasmatota, Halobacteria* and *Thermoprotei*, among others, where it is involved in C50 carotenoid biosynthesis (21). C50 carotenoids are derived from all-*trans*-lycopene (C40) (22), suggesting that the CrtDb subfamily emerged after the CrtN-P (C30) or CrtI-D (C40) subfamilies. In our controls, the location of CrtDb varies from basal to or intermediary between CrtI-N and CrtD-P subfamilies, showing the phylogenetic instability of this subfamily. However, the topology of the CrtI-N and CrtD-P clusters was stable in controls, except for some CrtP in *Firmicutes*, branching intermediary between CrtP and CrtD subfamilies (Fig. 6).

**Figure 6:**
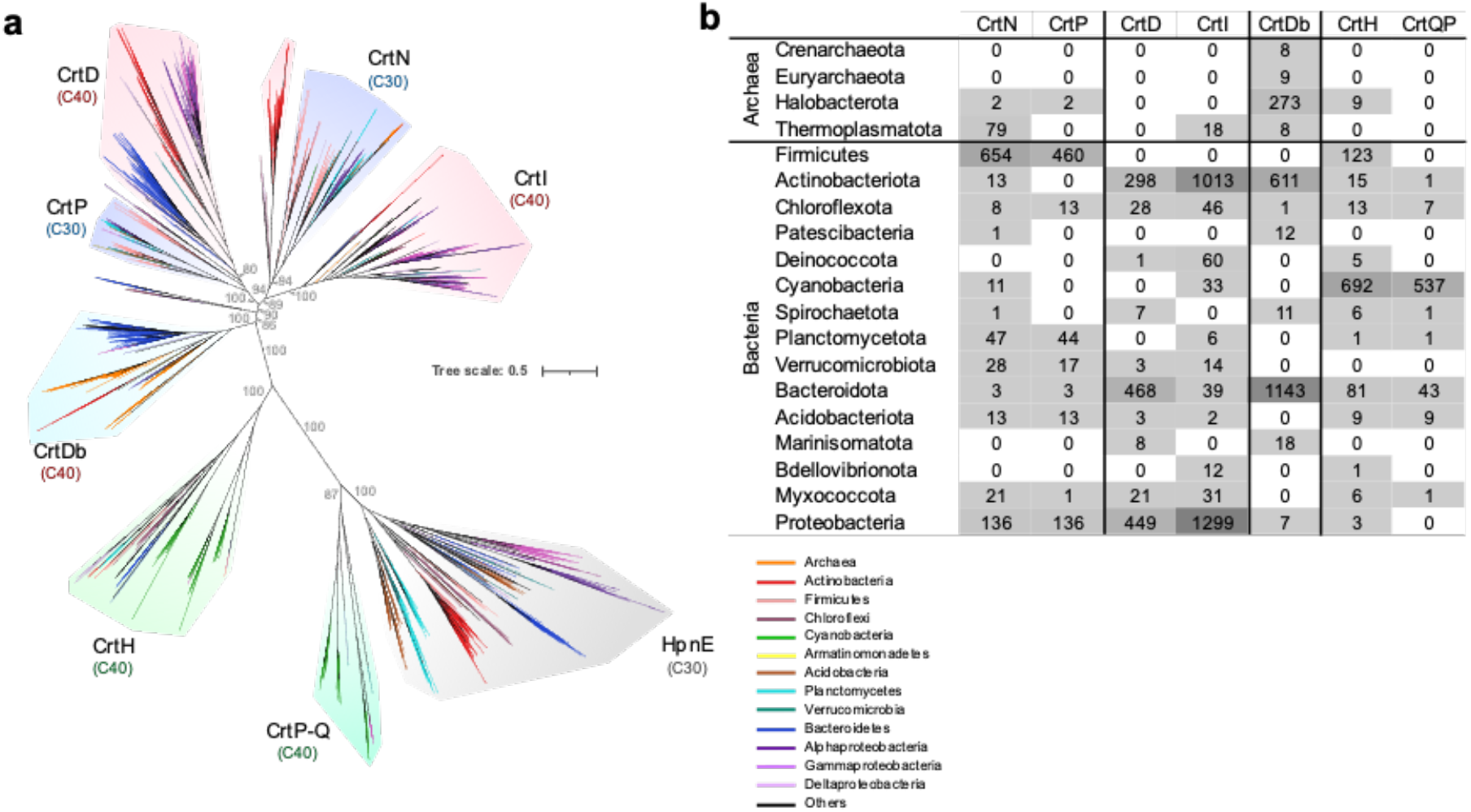
Phylogeny and taxonomic distribution of carotenoid amino oxidases. a) Maximum-likelihood reconstruction of selected non-redundant subfamilies of carotenoid amino oxidases. The subfamilies are highlighted and annotated according to the presence of characterized proteins. Branches are colored by taxonomic phyla. b) Taxonomic composition of the different subfamilies. Numbers indicates the number of sequences in each phylum. More information is shown in Data S1.

Enzymes of the C30 carotenoid pathway are closely related to those of the C40 carotenoid pathway of *Proteobacteria* (and others), but distantly related to those in cyanobacteria (Fig. 6). The presence of one Trans IPPS HH enzyme (CrtM or CrtB), two amino oxidases (CrtN-CrtP or CrtD-CrtI) and even a glucosyl-transferase (CrtQ or CrtX) in the C30 and C40 carotenoid operons, respectively, together with the fact that CrtN-CrtI and CrtP-CrtD branch together, respectively, suggests that these C30 and C40 pathways have a common origin, possibly due to neofunctionalization of an operon. In the phylogeny of carotenoid amino oxidases, CrtN and CrtP are branching closer to the deep nodes (Fig. 6), which suggests that the C30 carotenoid pathway diverged earlier than the C40 pathway. In agreement with this assumption, the direct evolution of the function of CrtB from CrtM has been demonstrated by random mutagenesis, but not the opposite (23). This assumption is further supported by the simpler molecular structure of C30 carotenoids, as they are generally formed by a linear and desaturated C30 backbone, which may also present one or two glucoses moieties at the acid extremes of the carotenoid molecule and a linear fatty acid radical linked to the sugar. By contrast, C40 carotenoids usually undergo cyclization at the extremes of the backbone, which add extra enzymatic steps involving the lycopene cyclases, the enzymes of the CrtY family. Together, these observations suggest that the C30 carotenoid pathway is ancestral to C40 (and C50) carotenoid biosynthesis and its most likely phylum of origin is *Firmicutes* or *Planctomycetes*, depending on whether it originated from 4,4’-diapophytoene or from squalene.

## Discussion

Here, we characterize the biosynthesis of C30 carotenoids via squalene. This molecule has traditionally been associated with the biosynthesis of polycyclic triterpenes. We demonstrate that it can also serve as an intermediate for C30 carotenoid biosynthesis and that this ‘squalene route’ to C30 carotenoid biosynthesis is widespread in *Bacteria*. The carotenoid profile of *P. limnophila* is characterized by an array of C30 carotenoids derived from 4,4-diapophytoene, in which unusual red carotenoids with carboxylic acid moieties predominate. This structural feature raises the possibility of new industrial and pharmaceutical applications for these molecules, since carotenoic acids have higher polarity and water solubility than common carotenoids.

Our results show that both carotenoids and polycyclic triterpenes have similar protective roles against environmental stresses such as desiccation and salinity. The resistance to stress was not associated with any particular molecule, demonstrating that C30 carotenoids and hopanoids are functionally related in *P. limnophila*. However, the possibility of complementary overproduction to compensate for a deficiency could not be discounted. In addition, these apparently similar roles of carotenoids and hopanoids could be influenced by remodeling other membrane features, such as the ratio of saturated:unsaturated fatty acids, as has previously been shown (8, 10). We also demonstrate that the accumulation of squalene in the *DcrtN*-Δ*shc* mutant is not toxic for the bacterium, but instead has a low, but apparent protective effect. Indeed, squalene can function by influencing membrane properties itself, as has been reported in *Halobacteria* (24), fungi (25) and mammals (26). These results provide further support for the related evolution between carotenoid and polycyclic triterpenes, and present squalene as a versatile compound in the evolution of triterpene derivatives.

The role of squalene as precursor of both carotenoids and polycyclic triterpenes raises important considerations for the diversification of these metabolites. C40 pathways have diversified more (CrtI-CrtD, CrtD-CrtI-50 or CrtP/Qc-CrtH) than the unique C30 carotenoid pathway. We infer that the C30 and C40 pathways (via CrtI-CrtD) have a common origin by neofunctionalization of an operon, that the C30 pathway most likely originated in *Firmicutes* or in *Planctomycetes*, and that it is ancestral to the CrtI-Db C40 pathway. We could also order the evolution of the pathways using the phylogenetic proximity between HpnE and CrtP-Qc. The HpnCDE enzymes show a more widespread and ancestral distribution in *Bacteria* (*i.e*. ancestral in phyla like *Actinobacteria*, *Planctomycetes* and *Proteobacteria*, among others^5^) than the CrtP-Q/H enzymes, which are present only in *Cyanobacteria* and *Chlorobi*, suggesting that squalene (HpnCDE) predates the cyanobacterial carotenoid pathway. Thus, the close relationship between the HpnE and CrtP-Qc subfamilies would imply that the cyanobacterial carotenoid pathway was derived from Hpn(CD)E-related enzymes. This is interesting, because *Cyanobacteria* shows the most ancestral association of Sqs with hopanoid biosynthesis. The related evolution of HpnE and CrtP-Qc (which raises the possibility that Trans IPPS HHs also co-evolved) supports the hypothesis that the origin of Sqs uncoupled polycyclic triterpenes from carotenoid biosynthesis, allowing the individualization of the pathways (4). Together, our results provide novel insights into the links between, and diversification of, carotenoid and polycyclic triterpene metabolisms, as well as optimizing the contextualization of these molecules, which are commonly used as biomarkers (27), throughout geological time scales. Our demonstration of the widespread occurrence of the squalene route to carotenoid biosynthesis increases the functional repertoire of squalene and establishes it as a general hub of polycyclic triterpenes and carotenoids biosynthesis.

## Methods

### Bacterial strains and culture conditions

The bacterial strains used in this work are listed in Table S1. *Escherichia coli* DH5α, used for cloning purposes, and *E. coli* SoluBL21 (28), used for carotenoid analysis, were grown in lysogeny broth (LB) medium at 37°C. *E. coli* SoluBL21 bearing the expression plasmids was grown in LB-based ZYM-5052 auto-induction medium(29). *Planctopirus limnophila* DSM3776^T^ was cultivated in M3 modified medium (30), pH 7.5, at 28°C. We added 1.5% bacto-agar to solid media for *E. coli* and 1% for *P. limnophila*. When required, kanamycin (Km) or gentamycin (Gm) was used at 50 and 20 μg mL^−1^ for *P. limnophila*. For *E. coli*, antibiotics were added at the following concentrations (μg mL^−1^): Km (25), Gm (10), chloramphenicol (Cl, 15), ampicillin (Ap, 100). Methyl viologen dichloride hydrate (paraquat, 98% purity) and Isopropil-β-D-1-tiogalactopiranósido (IPTG) were purchased from Merck.

### Plasmid construction

The oligonucleotides and plasmids used in this work are summarized in Tables S2 and S3, respectively. All DNA manipulations were performed using standard protocols. Plasmids used for gene deletion in a double event of homologous recombination were derived from pEX18Tc vector (31), which is a suicide plasmid containing a tetracycline resistance gene. To construct knockout plasmids for the target genes, fragments containing 700–1400 bp sequences of the flanking region of the target genes were amplified by PCR from genomic DNA of *P. limnophila* using the primers summarized in Table S2. The upstream and downstream fragments were cloned into pEX18Tc by three-way ligation using the appropriate restriction enzymes listed in Table S2. Finally, the Km or Gm resistance genes from the pUTminiTn5km (32) or pBBR1MCS-5 (33) plasmid, respectively, were subsequently cloned as a *BamHI* fragment between the flanking regions. Plasmids used for the reconstruction of carotenoid synthesis pathway in *E. coli* (Table S3) were divided into three categories: category 1, a plasmid that contains the genes required for the synthesis of the precursor IPP (Addgene); category 2, the plasmids necessary for squalene biosynthesis through squalene synthase (Sqs) or HpnCDE; and category 3, the plasmids that bear the genes required for carotenoid biosynthesis. All the genes contained in category 2 and 3 plasmids were amplified from genomic DNA using the pair of oligonucleotides listed in Table S2, cut with the appropriate restriction enzyme, and cloned into pBR322 (34) and pSEVA231 (35), respectively. The genes were expressed under the lacV5 promoter and a strong ribosome binding site. More details are presented in Table S3.

### Mutant strain construction

Genetic transformations of *P. limnophila* for the construction of deletion mutants were performed by electroporation, as previously described (30). In summary, fresh electrocompetent cells were prepared from 400 mL of a culture at OD_600_ 0.4 in modified M3. The cells were washed twice with ice-cold double-distilled sterile water (100 mL and then 50 mL) and once with 2 mL of ice-cold 10% glycerol. The pellet was then resuspended in 400 μL of ice-cold 10% glycerol, and aliquots of 100 μL were dispensed into 0.1 mm gapped electroporation cuvettes along with 1 μg of plasmid DNA and 1 μL of Type-One restriction inhibitor (Epicentre). Electroporation was performed with a Bio-Rad Micropulser (Ec3 pulse, voltage [V] 3.0 kV). Electroporated cells were immediately recovered in 1 mL of cold fresh medium and incubated at 28°C for 2 h with shaking. The cells were then plated onto agar plates supplemented with Km or Gm and were incubated at 28°C until colony formation after approximately 7 days. Colonies were segregated onto fresh selection plates and genotyped by PCR and sequencing.

For random mutagenesis by transposition, 1 μL of EZ-Tn5 solution and 1 mL of Type-One restriction inhibitor were electroporated following the aforementioned protocol. The cells were then plated onto modified M3 supplemented with Km and incubated at 28°C until colony formation. White colonies were segregated onto fresh selection plates. To verify Tn5 insertions and their locations, DNA was isolated using the Wizard Genomic DNA Purification Kit (Promega), and analyzed by semirandom PCR (36). Genomic DNA was used as the template DNA in a 20 μL PCR mixture containing primer Map Tn5 A fwd and a mix of primers CEKG 2A, CEKG 2B and CEKG 2C; 1 μL of a 1:5 dilution of this reaction mixture was used as the template DNA for a second PCR performed with primers Map Tn5 B fwd and CEKG 4. For the first reaction, the thermocycler conditions were 94°C for 2 min; followed by six cycles of 94°C for 30 s, 42°C for 30 s (with the temperature reduced 1°C per cycle), and 72°C for 3 min; and then 25 cycles of 94°C for 30 s, 65°C for 30 s, and 72°C for 3 min. For the second reaction, the thermocycler conditions were 30 cycles of 94°C for 30 s, 65°C for 30 s, and 72°C for 3 min. The DNA of purified PCR products (GFX PCR DNA and Gel Band Purification Kit GE Healthcare) was sequenced using primer Map Tn5 B fwd.

### Carotenoid pathway reconstruction

To construct the different *E. coli* expression strains, the appropriate plasmids (Table S3) were transformed into *E. coli* SoluBL21 by heat shock. SoluBL21 competent cells were prepared by TSS methods **(37)**. Cells were plated in LB containing Cl, Ap and Km. When liquid cultures were required, preinocula with the appropriate antibiotics were grown in LB at 37°C until saturation. Once grown, the cultures were diluted to OD_600_ 0.1 in fresh media and then grown at 37°C until OD_600_ 0.4, at which point they were induced with IPTG 0.5 mM (Merck) and incubated at 28°C for 48 h. Cell were collected by centrifugation at 6,000 xg and 4°C and the pellets were kept at −80°C.

### Carotenoid production and extraction

For pigment production, strain cultures (250 to 1000 mL flasks) were performed according to respective *P. limnophila* and *E. coli* culture conditions. Cells were then harvested by centrifugation at 5,000 xg and 4°C, and the pellets were washed with phosphate buffer 1X (6.05 g L^−1^ of Na_2_HPO_4_.12H_2_O and 1.0 g L^−1^ of KH_2_PO_4_), frozen at −80°C, and lyophilized (VirTis BenchTop 2 K Freeze Dryer, SP Industries Inc.). Approximately 0.15 g of lyophilized biomass was sequentially extracted with ethanol, methanol and acetone (5 mL each) until no more color was extracted. Extraction was aided by vortex shaking for 1 min and sonication for 30 s. Extraction fractions were collected after centrifugation of samples (5,000 xg at 4°C), the solvent was evaporated to dryness under vacuum in a rotary evaporator (<30°C), and the dry extract was dissolved in acetone-ethanol (1:1) for chromatographic analysis. We performed all the operations under dimmed light to avoid isomerization and photo-degradation of carotenoid pigments.

### Carotenoid identification

Carotenoid identification was based on the chromatographic and UV-visible spectroscopic (UV-visible and mass spectrometry) data obtained by HPLC coupled with a diode array detector (HPLC-DAD) and HPLC coupled with a mass spectrometer (HPLC-MS(APCI)). Data was compared with those of literature values (12, 18, 38–43). HPLC-DAD analysis was carried out using a Waters e2695 Alliance chromatograph fitted with a Waters 2998 photodiode array detector and controlled with Empower2 software (Waters Cromatografía, SA, Barcelona, Spain). The separation was performed in a reverse-phase C18 (20 mm x 4.6 mm i.d., 3 μm, Mediterranea SEA18; Teknokroma, Barcelona, Spain) fitted with a guard column of the same material (10 mm x 4.6 mm). The chromatographic method used was previously described in Delgado-Pelayo *et al*. (44), although we added formic acid (0.1 % final concentration) to the mobile phase. Briefly, carotenoid separation was carried out by a binary-gradient elution using an initial composition of 75 % acetone and 25 % deionized water (both containing 0.1 % formic acid), which was increased linearly to 95 % acetone in 10 min, held for 7 min, raised to 100 % in 3 min, and held for 10 min. Initial conditions were reached in 5 min. The temperature of the column was kept at 25°C and the sample compartment was refrigerated at 15°C. An injection volume of 10 μL and a flow rate of 1 mL min^−1^ were used. Detection was performed at 500 nm for major pigments and 370 nm for early precursors, and the online spectra were acquired in the 350–700 nm wavelength range. HPLC-MS(APCI) analysis was carried out with a Dionex Ultimate 3000RS U-HPLC (Thermo Fisher Scientific, Waltham, MA, USA) coupled in series with a diode array detector (DAD) and a micrOTOF-QII high resolution time-of-flight mass spectrometer (UHR-TOF) with qQ-TOF geometry (Bruker Daltonics, Bremen, Germany) and fitted with an APCI (atmospheric pressure chemical ionization) source. The chromatographic conditions were identical to those described for HPLC-DAD analysis. A flow-split of the eluent from the DAD detector was set up in order to allow a 0.4 mL/min flow rate directly into the mass spectrometer (connected in series after the DAD detector). The instrument control was performed using Bruker Daltonics Hystar 3.2 software and data evaluation was performed with the Bruker Daltonics DataAnalysis 4.0 software. The MS parameters were set as follows: positive mode; current corona, 4000 nA; source (vaporizer) temperature, 350°C; drying gas, N2; gas temperature, 250°C; gas flow, 4 L/min; nebulizer pressure, 60 psi; scan range of m/z 50–1200.

For alkaline hydrolysis of carotenoid extracts from *P. limnophila* wild type, 1 mL of crude extract was evaporated to dryness under a nitrogen stream, and the residue was dissolved in 3 mL of 0.25 N NaOH (aqueous) and left to react for 24 h at room temperature (<25°C) in the dark. The mixture was acidified with formic acid and the pigments recovered with diethyl ether. The ether phase was collected, evaporated under nitrogen stream and dissolved in acetone-ethanol (1:1) for chromatographic analysis.

### Analysis of squalene by gas chromatography

Lyophilized biomass pellet (0.1 g) was submitted to alkaline hydrolysis with 2 mL of 2 % (w/v) KOH-ethanol at 80°C for 15 min. Squalane (20 μL; stock solution 10.8 mg mL^−1^) was added as internal standard. After cooling to room temperature, the mixture was diluted with 3 mL of distilled water and extracted with 1 mL n-hexane. An aliquot of the upper hexane phase (0.5 mL) was transferred to a vial for GC-FID analysis. Gas chromatography analysis were performed on an Agilent Technologies 7890A gas chromatograph (Agilent Technologies España, S.L., Madrid, Spain) fitted with a flame ionization detector, a split/splitless injector, and a 7683B series automatic liquid sampler. The chromatograph was fitted with a HP-5 capillary column (J&W Scientific; 30 m length; 0.32 mm i.d.; 0.25 μm thickness). Helium was used as carrier gas with a constant linear flow of 1.75 mL min^−1^. The injector and detector temperatures were 300°C and 325°C, respectively. The oven temperature started at 250°C and increased at a rate of 4°C min^−1^ to 270°C, where it was held for 3 min. The injection volume was 1 μL at a split ratio of 1:20.

### Phenotypic stress analysis

For the physiological assays, preinocula of the *P. limnophila* wild type and mutants were grown in liquid media with antibiotics until saturation. Stress assays carried out in solid media (1% agar) used 5 to 10 μL drops from ten-fold serial dilution (cultures were adjusted to the same OD_600_). For the oxidative stress assays, cells were grown in paraquat-supplemented solid medium to a final media concentration of 2 μM. To evaluate the resistance to osmotic stress, we grew cells on plates supplemented with 15 mM sodium chloride. For desiccation assays, cells were placed on a nitrocellulose membrane and left to air-dry in a laminar flow cabinet for 1 h (control was left for 5 min to allow the drops to dry). Then, the membranes were placed onto solid medium and incubated. For the temperature assays, saturated cultures were further diluted to OD_600_ 0.03 and grown at different temperatures (16, 22, 28, 32, 36, 38 and 40°C) until cultures reached stationary phase. OD_600_ measurements of the cultures were taken regularly along the growth curve. A temperature stress assay was also performed by subjecting each strain to three freeze-thaw cycles. Aliquots of cell suspensions grown at 28°C were frozen at −20°C. After 24, 48, and 72 h, cells were thawed, and the viability was determined in solid media by plating 10 μL drops from ten-fold serial dilution and compared with the viability of non-treated cells. The remaining sample volume was re-frozen for subsequent freeze-thaw cycles. Statistical analysis was performed using IBM SPSS version 25; SPSS Science (Chicago, IL, USA). Student’s t-test was used to compare mean growth rates. A significance level of P < 0.05 for the 95% confidence interval was chosen to define the statistical significance.

### Phylogeny

To infer the evolution of carotenoid amino oxidases, we searched for homologous sequences to amino oxidases such as HpnE, CrtN, and CrtI. We performed PHMMER searches (45) with an e-value threshold of 1e^−5^, against a local database containing NCBI proteomes of those organisms described in GTDB (version 120 (46)). We combined the resulting target sequences (~14,000) into a single dataset, aligned them using MAFFT(47), and trimmed gap positions using trimAL (48) and some other non-informative regions manually. With this alignment, we then performed a guide tree using Fasttree (default parameters) (49), and selected the subfamilies of interest according to the presence of characterized enzymes. For each subfamily, we removed non-redundant sequence for taxonomic classes by applying different cut-offs depending on the number of sequences (20 sequences, 95 % identity; up to 250 sequences, 55 %). These individual reduced subfamilies were again combined to performed the final phylogenies of carotenoid/squalene amino oxidases and those for CrtN/P. We aligned these datasets using MAFFT-linsi, trimmed gap positions and removed spurious sequences. These final alignments were used for phylogenetic reconstructions using IQ-TREE (50). We obtained branch supports with the ultrafast bootstrap (51), and the evolutionary models of each set of sequences were automatically selected using ModelFinder (52) and chosen according to BIC criterion. All trees were visualized and annotated using iTOL (53).

For the phylogenetic profiles mapped onto the CrtN/P phylogeny, we made use of a previously defined dataset for HpnCDE, CrtB/M, Sqs and Shc subfamilies (4).

### Genome context

We defined the genome context as the arrangement of neighboring genes relative to the gene of interest. To analyze the genome contexts of genes containing the amino oxidase domain (PF01593), we extracted the genomic sequence of the genes 10 Kb upstream and downstream. We extracted the coding DNA sequence from these genomic fragments using PRODIGAL (54), annotated the coding proteins using the PFAM database (55) running HMMSCAN (45), and parsed the output to keep the longest coverage and best e-value in order to minimize the effect of overlapping domains. To identify the genes that are in the genome context of *crtN* (Data S2), we took all the coding genes containing the amino oxidase and SQS PSY domains, and searched against a homemade database of the different Trans IPPS HH and amino oxidases subfamilies using PHMMER with 1e^−20^ as the e-value threshold.

## Author contributions

C.S.M, E.R.M and D.P.D designed the study. C.S.M performed *in silico* analysis. V.H. and E.R.M constructed the mutants and performed the physiological assays. D.H. chemically characterized *P. limnophila* triterpenoids. All authors analyzed and interpreted data, and contributed to writing the manuscript.

## Data and materials availability

All data needed to evaluate the conclusions in the paper are present in the paper and/or the Supplementary Materials.

## Supplementary Materials

**Fig. S1.**
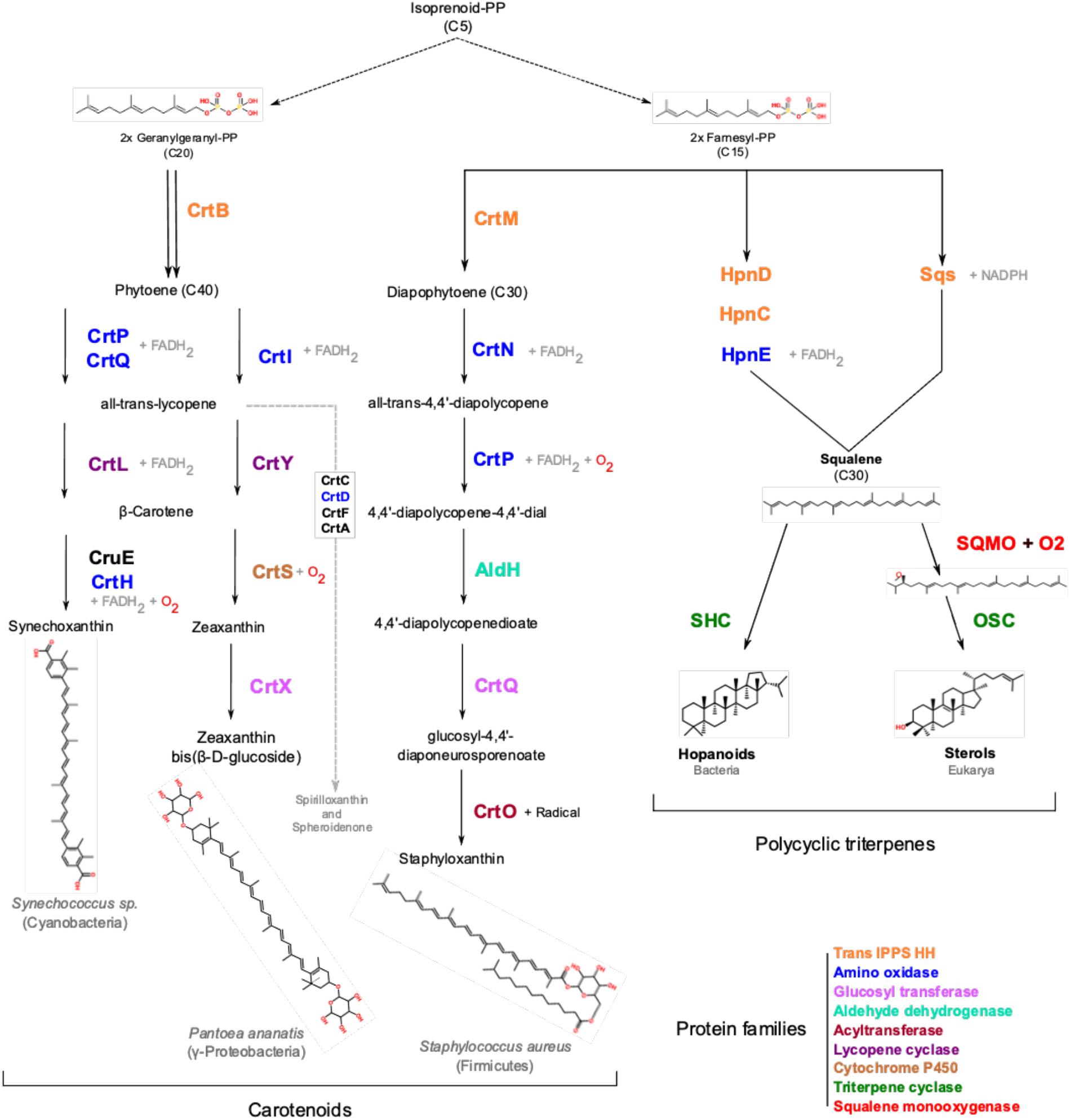
Schematic of the canonical polycyclic and linear terpenoid biosynthesis pathways. Biosynthetic pathways of polycyclic triterpenes and carotenoids (C30 and C40), starting from farnesyl-PP or geranylgeranyl-PP. Homologous enzymes are shown in the same color to illustrate the evolutionary relationship and homologies between the pathways. Note that the pathways shown are representative; alternative routes are possible.

**Fig. S2.**
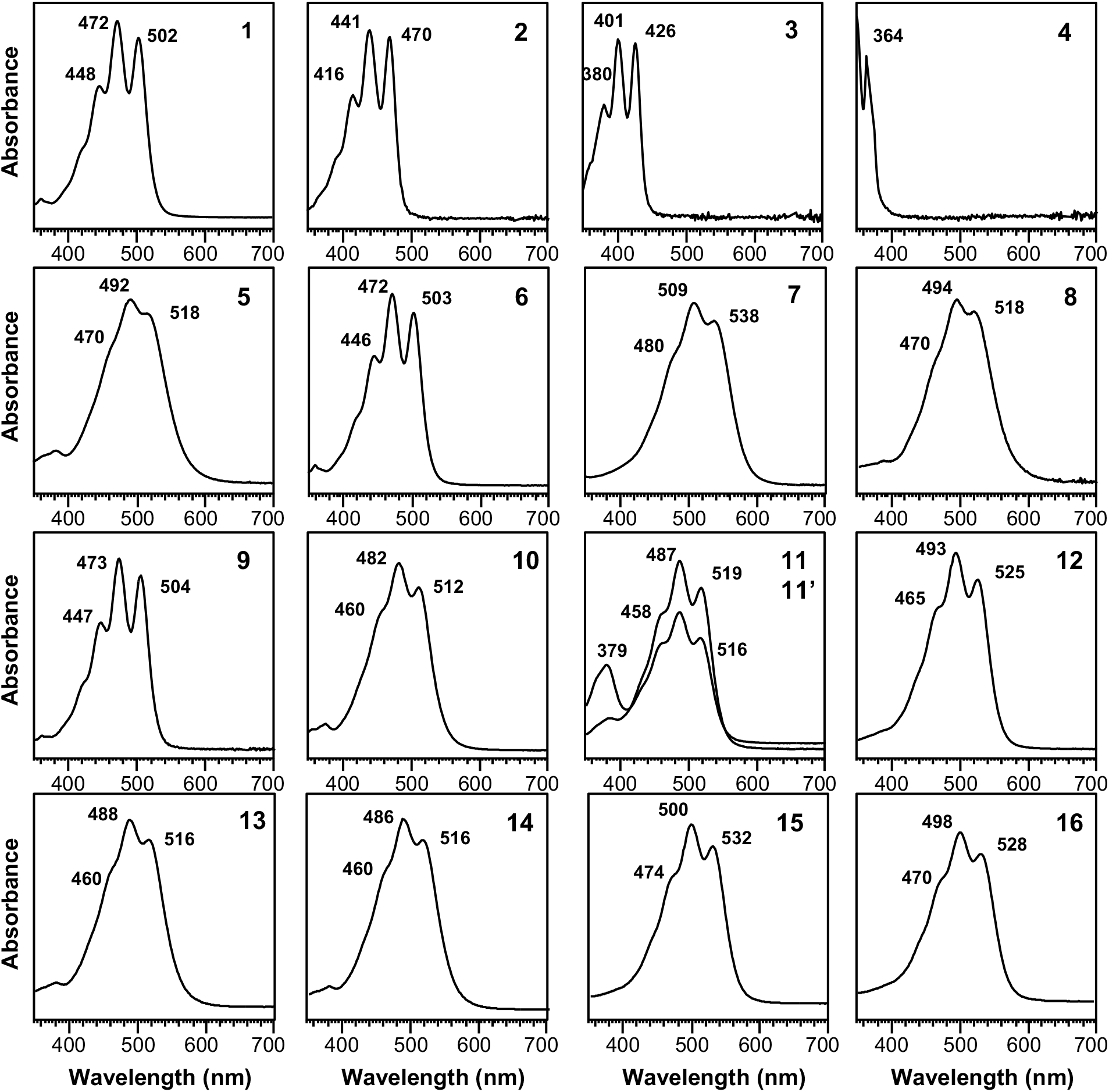
UV-visible spectra of the carotenoids from *P. limnophila* mutants. Spectrum numbers are in accordance with chromatogram peaks in Figure 2 and compounds in the pathway scheme (Fig. 3): 4,4’-diapolycopene (1); 4,4’-diaponeurosporene (2); 4,4’-diapo-ζ-carotene (3); 4,4’-diapophytofluene (4); 4,4’-diapolycopen-4-al (5); 4,4’-diapolycopene-4-ol (6); 4,4’-diapolycopene dial (7); 4,4’-diapolycopene-4-ol-4’-al (8); 4,4’-diapolycopene-4,4’-diol (9); 4,4’-diapolycopenoic acid (10); 4,4’-diapolycopen-4’-al-4-oic acid (11 & 11’); 4,4’-diapolycopen-4,4’-dioic acid (12); glycosyl esters of 4,4’-diapolycopenoic acid (13 & 14); and glycosyl esters of 4,4’-diapolycopen-4,4’-dioic acid (15 & 16).

**Fig. S3.**
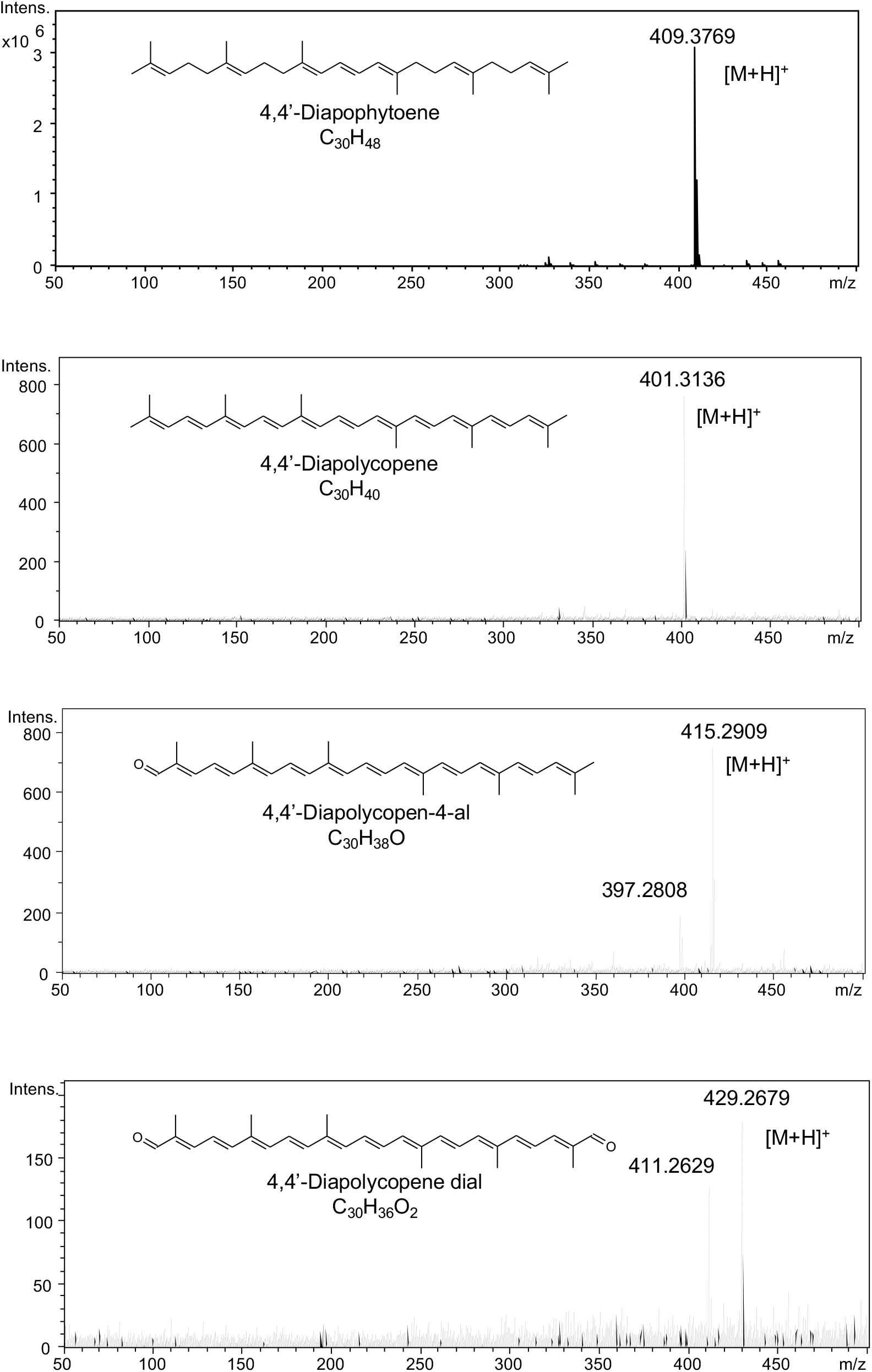
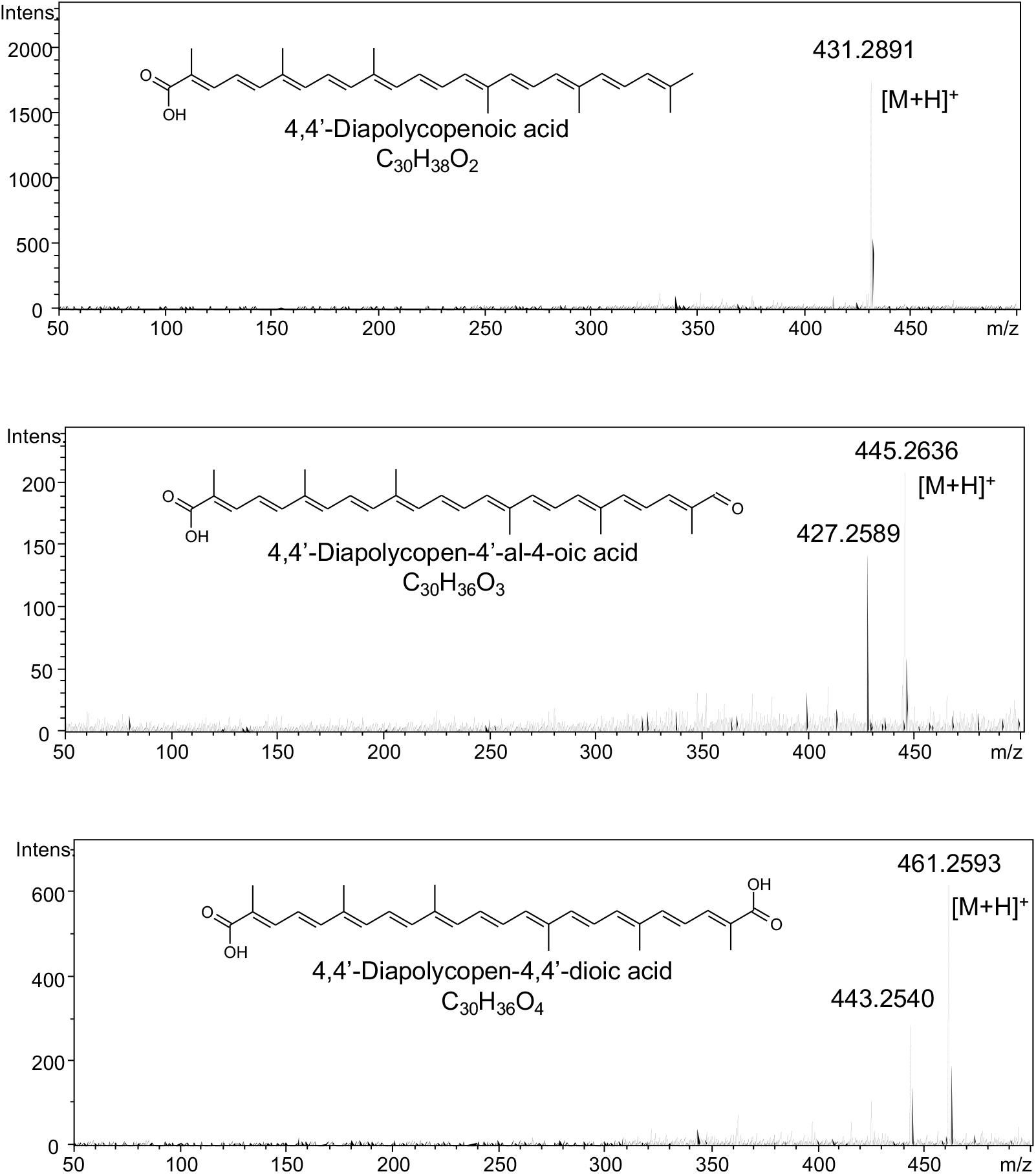
Mass spectra obtained by HPLC-MS(APCI) for the major C30 carotenoids produced by *P. limnophila*.

**Fig. S4.**
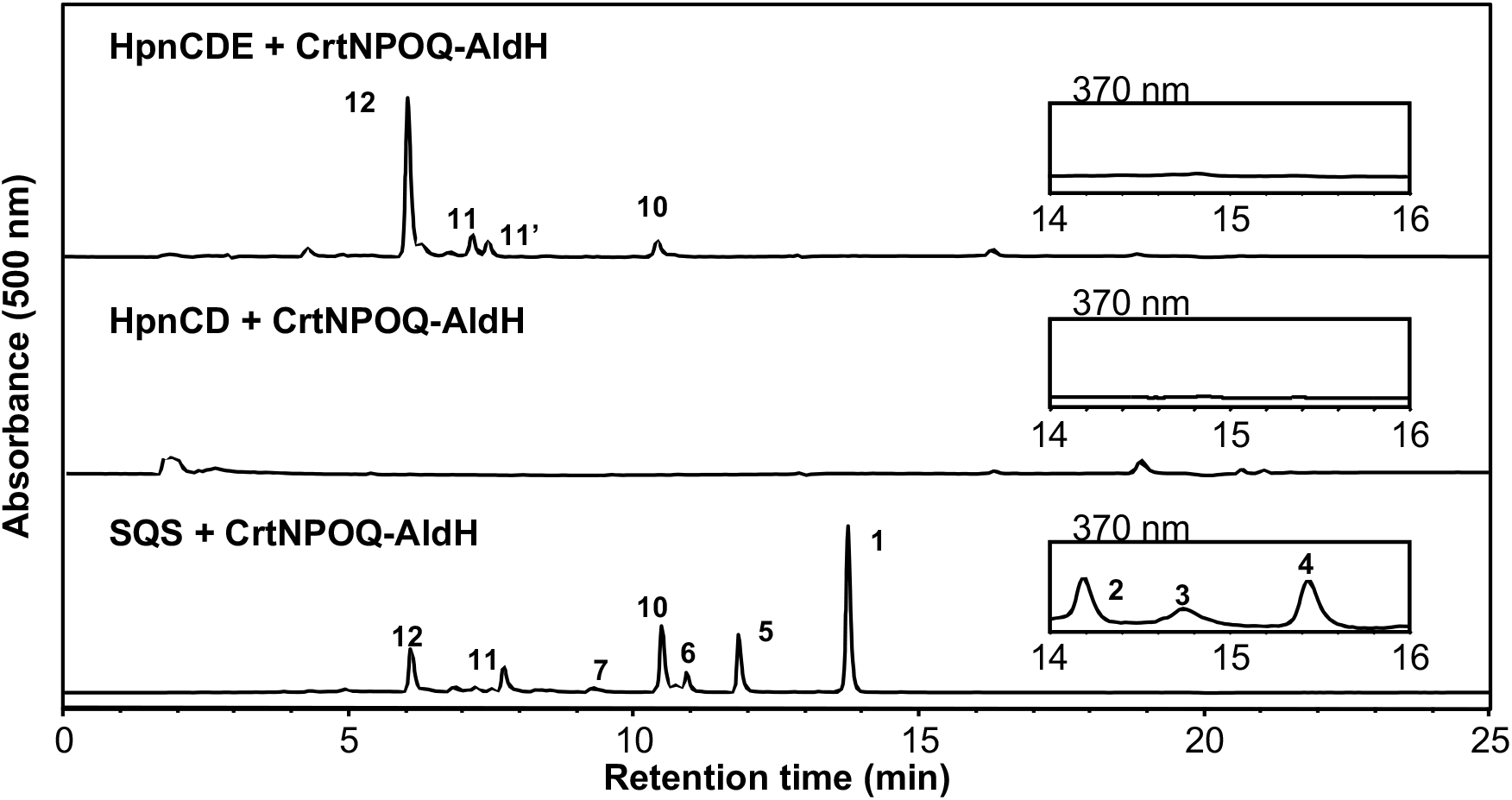
Analysis of the carotenoid heterologous expression system. HPLC chromatogram corresponding to the carotenoid analysis of *E. coli* recombinants bearing plasmid 1 (IPP supplier), a plasmid producing squalene (via HpnCDE or SQS) or its precursor (via HpnCD), and a plasmid containing *crtNPOQ-aldH* genes. Detection wavelengths at 370 and 500 nm. Peaks numbers as in Figure 2 and pathway scheme (Fig. 3). Peaks: 4,4’-diapolycopene (1); 4,4’-diaponeurosporene (2); 4,4’-diapo-ζ-carotene (3); 4,4’-diapophytofluene (4); 4,4’-diapolycopen-4-al (5); 4,4’-diapolycopene-4-ol (6); 4,4’-diapolycopene dial (7); 4,4’-diapolycopenoic acid (10); 4,4’-diapolycopen-4’-al-4-oic acid (11 & 11’); 4,4’-diapolycopen-4,4’-dioic acid (12).

**Fig. S5.**
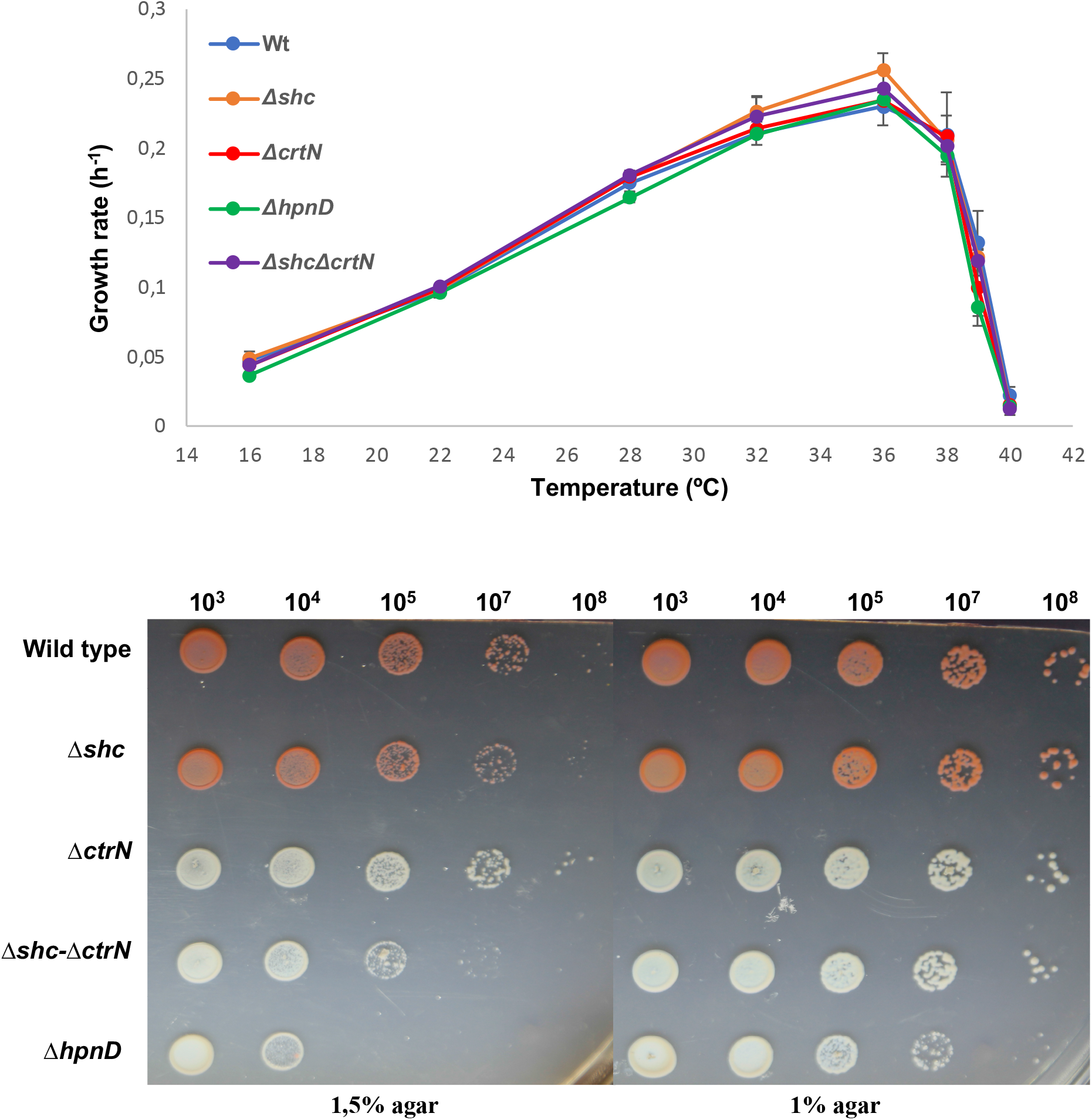
Phenotypic assays of selected *P. limnophila* mutants. a) growth rate at different temperatures (16-40°C) and b) growth in solid medium (1.5 % vs 1% agar).

**Fig. S6.**
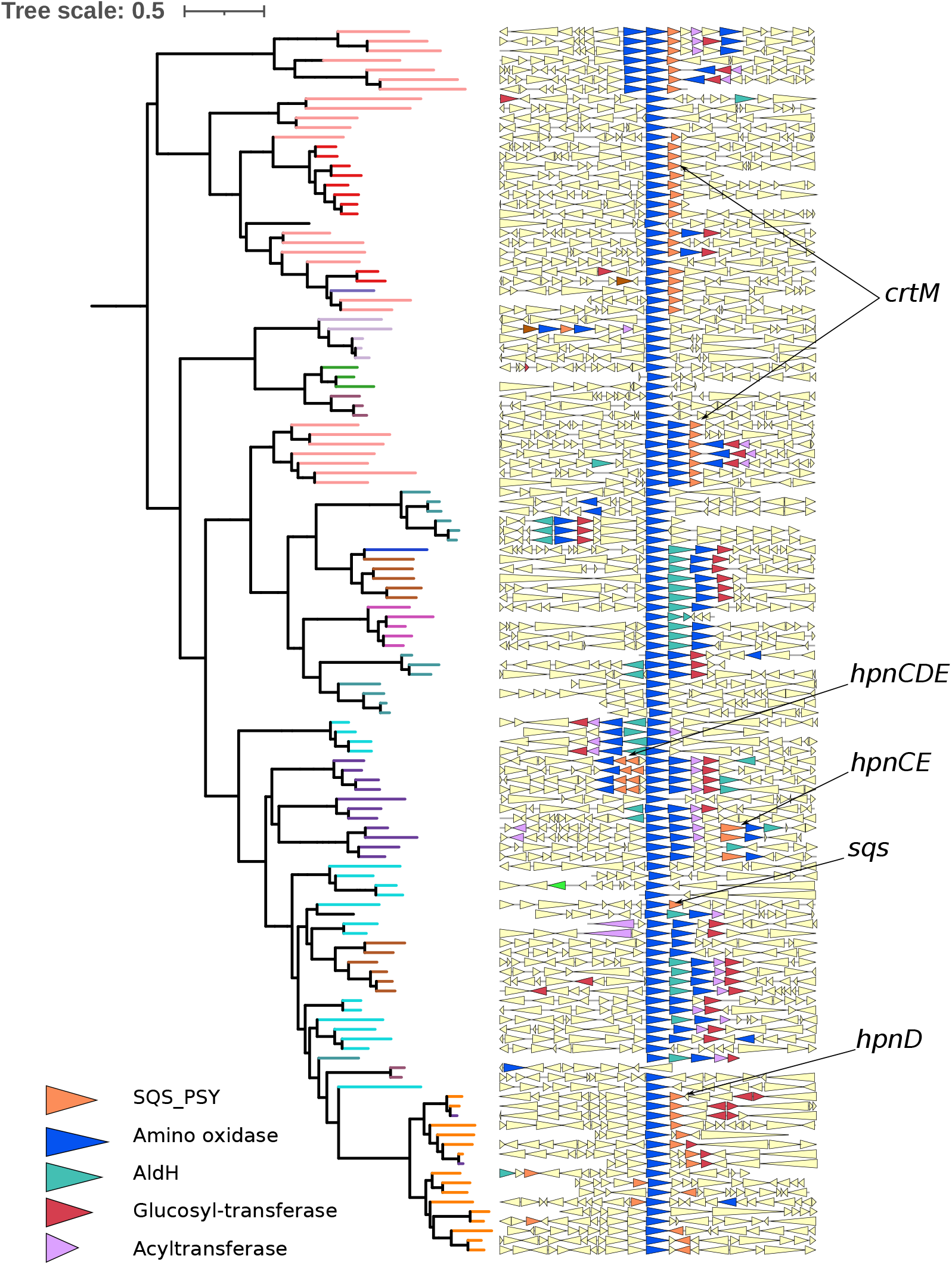
Genomic context of *crtN* genes across prokaryotes. Pruned tree of CrtN phylogeny and synteny of *crtN* gene across 10 Kb upstream and downstream. Colored genes contain Pfam domain of interest. The cooccurrence of possible sources of precursors in the genomic context of *crtN* are indicated (*crtM, hpnCDE* or *sqs*).

**Fig. S7.**
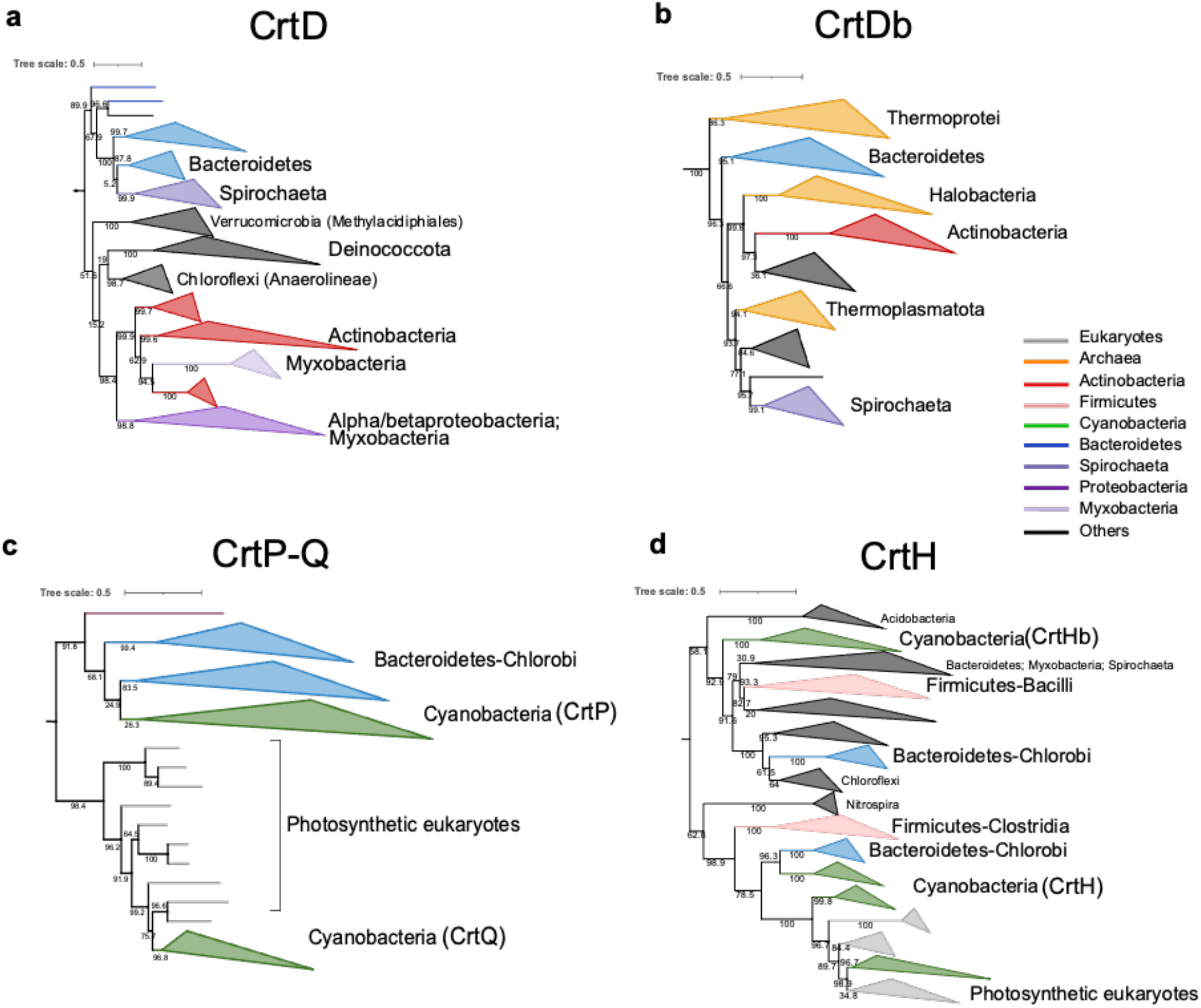
Pruned subfamilies from the phylogeny of carotenoid amino oxidases. a) CrtD b) CrtDb involved in the C50 pathway, c) CrtP-Q, and d) CrtH of the cyanobacterial pathway. Minor groups are indicated with collapsed branches in dark gray.

**Table S1.**
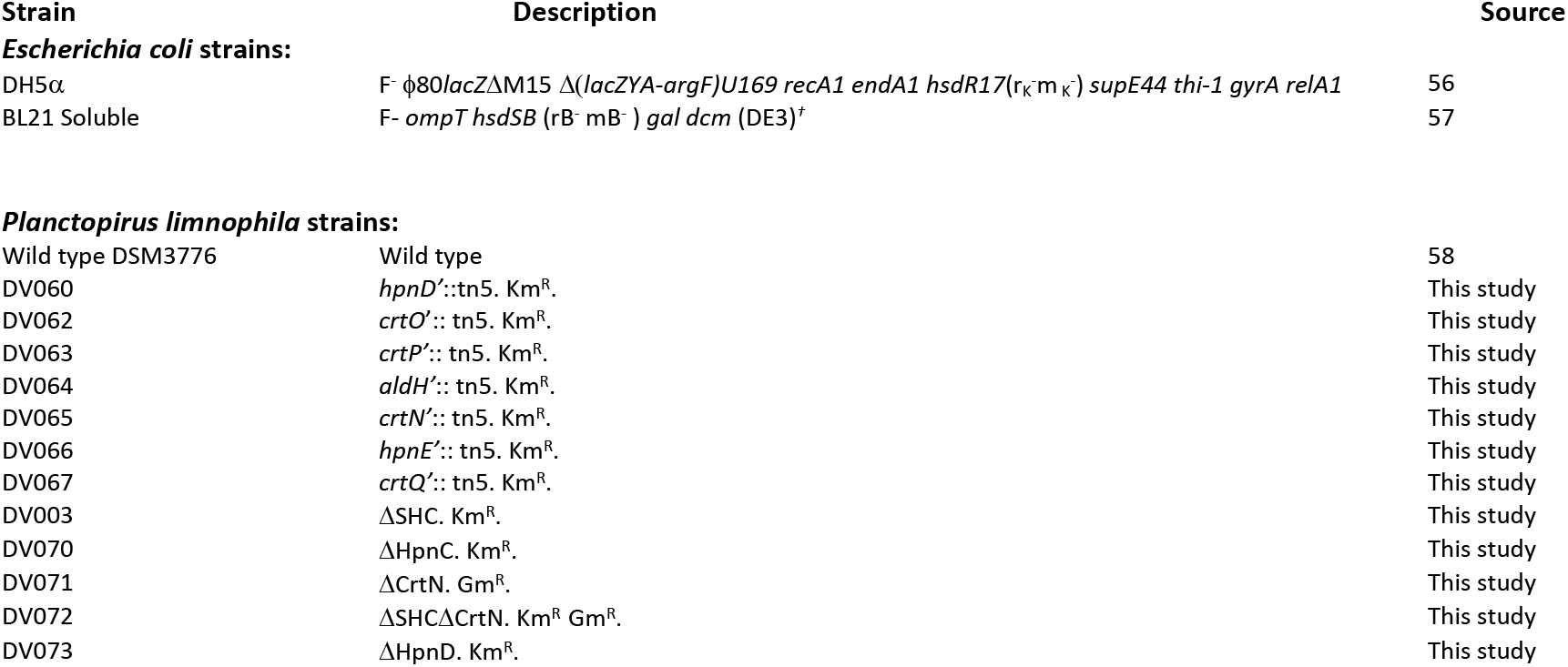
Strains used in this work.

**Table S2.**
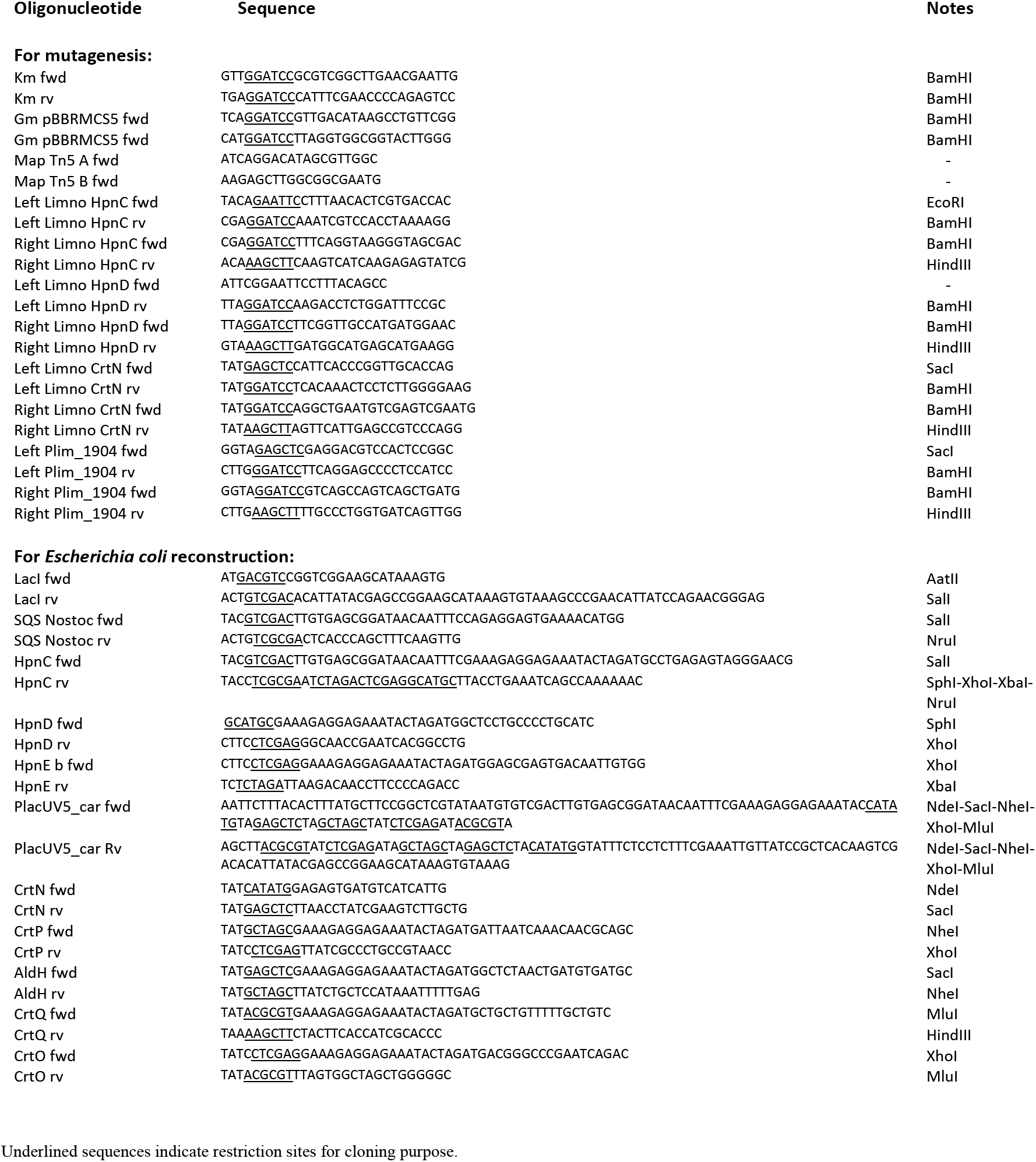
Oligonucleotides used in this work.

**Table S3.** Plasmids used in this work.

| Plasmid | Description | Source |
| --- | --- | --- |
| **For *P. limnophila* mutagenesis:** |  |  |
| pEX18Tc | Suicide plasmid. SacB.Tet<sup>R</sup>. | 59 |
| pDV008 | 1065 bp upstream and 1346 bp downstream of squalene hopene cyclase (*shc*) gene from *P. limnophila* flanking a kanamycin resistance gene cloned into pEX18Tc. Km<sup>r</sup>, Tc<sup>r</sup>. | This study |
| pDV103 | 859 bp upstream and 827 bp downstream of *hpnC* gene from *P. limnophila* flanking a kanamycin resistance gene cloned into pEX18Tc. Km<sup>r</sup>, Tc<sup>r</sup>. | This study |
| pDV111 | 837 bp upstream and 838 bp downstream of *crtN* gene from *P. limnophila* flanking a kanamycin resistance gene cloned into pEX18Tc. Gm<sup>r</sup>, Tc<sup>r</sup>. | This study |
| pDV118 | 776 bp upstream and 777 bp downstream of *hpnD* gene from *P. limnophila* flanking a kanamycin resistance gene cloned into pEX18Tc. Km<sup>r</sup>, Tc<sup>r</sup>. | This study |
| **For *E. coli* reconstruction:** |  |  |
| **Category 1:** |  |  |
| pBbA5c-MevT(CO)<br>MBis (CO, ispA) | p15A ori, AtoB(co)-HMGS(co)-HMGR(co)-MK(co)-PMK(co)-PMD-Idi-IspA. Cm<sup>R</sup>. | Addgene |
| **Category 2:** |  |  |
| pBR322 | pBR322 ori. Amp<sup>R</sup> Tet<sup>R</sup>. | 60 |
| pDV105 | pBR322 ori, lacUV5 promoter, *lacI<sup>q</sup>*. Amp<sup>R</sup>. | This study |
| pDV106 | *Anabaena* sp. PCC 7120 squalene synthase (SQS) expression plasmid. Derived from pDV105. | This study |
| pDV120 | *P. limnophila* HpnCDE expression plasmid. Codon optimize for *Escherichia coli*. Derived from pDV105. | This study |
| pDV122 | *P. limnophila* HpnCD expression plasmid. Codon optimize for *E. coli*. Derived from pDV105. | This study |
| **Category 3:** |  |  |
| pSEVA231 | pBBR1 ori, Km<sup>R</sup>. | 61 |
| pDV112 | pBBR ori, lacUV5 promoter, Km<sup>R</sup>. | This study |
| pDV113 | *P. limnophila* CrtN expression plasmid. Derived from pDV112. | This study |
| pDV115 | *P. limnophila* CrtN-CrtP expression plasmid. Derived from pMPO112. | This study |
| pDV114 | *P. limnophila* CrtN-CrtP-AldH expression plasmid. Derived from pDV112. | This study |
| pDV123 | *P. limnophila* CrtN-CrtP-AldH-CrtQ expression plasmid. Derived from pDV112. | This study |
| pDV117 | *P. limnophila* CrtN-CrtP-AldH-CrtQ-CrtO expression plasmid. Derived from pDV112. | This study |

**Data S1. (separate file)**

**Distribution of the main enzymes of this analysis.** Distribution of the main enzymes involved in polycyclic triterpene and carotenoid biosynthetic pathways across bacteria, archaea and some eukaryotes. The names of the phyla and classes are according to GTDB taxonomy and are sorted alphabetically by phylum. Numbers indicate the number of genomes with at least one respective gene.

**Data S2. (separate file)**

**Copy number of the main enzymes of this analysis.** Presence and number of copies of carotenoid amino oxidases and Trans IPPS HHs in the genomic context of *crtN* across the analyzed bacteria. Numbers indicate the number of copies in the genomic context of each subfamily. Light gray denotes low number of gene copies, and dark gray denotes higher number of gene copies in the respective genome contexts.

